# The Extracellular Matrix Regulates Tissue Mechanics to Enable Cyclic Lymph Node Remodelling for Sustained Immunity

**DOI:** 10.64898/2026.08.17.745227

**Authors:** Veronika Lachina, Pablo Vicente-Munuera, Amy Llewellyn, Spyridon Makris, Agnesska C. Benjamin, Kalnisha Naidoo, Yanlan Mao, Sophie E. Acton

**Affiliations:** Stromal Immunology Group, Laboratory for Molecular Cell Biology, University College London, Gower Street, London, WC1E 6BT, United Kingdom; Tissue Mechanics Lab, Laboratory for Molecular Cell Biology, University College London, Gower Street, London, WC1E 6BT, United Kingdom; Translational Pathology, Department of Physiology, King’s College London, London, WC2R 2LS, United Kingdom

**Keywords:** lymph nodes, immunology, biomechanics, immune response, extracellular matrix, tissue mechanics

## Abstract

Tissue shape and function are defined by the mechanical interactions of cellular and extracellular components. Lymph nodes cyclically remodel in response to immune challenges whilst preserving essential tissue architecture, comprised of a stromal fibroblastic reticular cell (FRC) network and extracellular matrix (ECM) it ensheaths. We experimentally quantified the contribution of ECM to the viscoelastic properties of the FRC network to parameterise an *in silico* model exploring factors affecting the FRC network’s adaptation to lymph node expansion. The balance between tissue pressure, FRC contractility, and ECM stiffness permit robust tissue remodelling, while maintaining physiological geometries. Local perturbation of ECM stiffness or FRC contractility disrupts force distribution and impacts FRC proliferation and tissue expansion. Spatially dispersed perturbations exert higher impact on tissue remodelling than equivalent localised perturbations, with effects propagating across the network. The lymph node provides a system for studying the integration of cellular and extracellular mechanics during dramatic tissue remodelling.

## Introduction

Few biological tissues undergo such frequent and extensive changes in size as lymph nodes. As secondary lymphoid organs, lymph nodes coordinate adaptive immune responses. During immune activation, they reversibly expand two-to ten-fold before returning to near-homeostatic size as the immune response resolves^1–3^. The magnitude of lymph node expansion varies between immunising agents and predicts immune response efficacy, suggesting that expansion is a determinant rather than merely a consequence of effective immunity^4^. The highly organised lymph node architecture, required for effective immune responses, must be preserved throughout expansion. Lymph nodes are enclosed by a fibrous capsule and are organised into T- and B-cell compartments interconnected by stromal, vascular, and extracellular matrix networks. Remarkably, repeated, large-scale lymph node deformation occurs tens to hundreds of times throughout life without mechanical failure. How lymph nodes achieve this remarkable mechanical robustness remains unknown. Here, we ask what mechanisms enable cyclic yet controlled lymph node expansion to occur without tissue failure.

Mechanical robustness depends on the regulation of forces acting within the tissue^5^. Lymph node tissue size emerges from the balance between contractile forces of the fibroblastic reticular cell (FRC) network and the pressure generated by infiltrating and proliferating lymphocytes^2,3,6^. During an immune response, dendritic cells expressing C-type lectin 2 (CLEC-2) engage podoplanin (PDPN) on FRCs, transiently reducing actomyosin contractility within the reticular network^2,3,7^. Concurrently, pressure in the tissue increases as lymphocytes become retained within the lymph node owing to sphingosine-1-phosphate receptor 1 (S1P1) downregulation^1–3^. Together, these shifting forces drive tissue expansion through stretching of the reticular network^2,3^. Yet it remains unknown how forces generated at the cellular scale are coordinated across the tissue to enable robust organ-scale expansion while preserving lymph node architecture and FRC network’s tightly regulated geometry and topology.

FRCs deposit and ensheathe extracellular matrix (ECM) conduits consisting of a fibrillar collagen core surrounded by a basement membrane^8,9^. In many tissues, the ECM is considered the principal force-bearing component responsible for maintaining tissue mechanical integrity^10^. However, how tissue architecture, mechanical properties, and function collectively emerge from dynamic interactions between cells and the extracellular matrix is unclear. Lymph nodes provide a unique system in which to address this interaction. In the lymph node, FRCs and ECM are mechanically integrated, and their relative mechanical contributions can be interrogated during cycles of tissue remodelling. Although considerable progress has been made in quantifying the mechanical forces present in the lymph node^2,3^, experimentally disentangling the relative contributions of cellular versus non-cellular components remains challenging. We thus developed a mechanical vertex model of the lymph node FRC network that incorporates lymphocyte-derived pressure, FRC actomyosin contractility, and ECM elasticity. Comparing live tissue experimental data with *in silico* perturbations, we identify ECM as a key contributing factor to tissue elasticity and force buffering during lymph node homeostasis and expansion.

We further used our model to investigate how pathological alterations in FRC contractility and ECM mechanics influence lymph node expansion and tissue robustness. Chronic inflammatory conditions can lead to persistent ECM accumulation and the emergence of hypercontractile FRCs, fundamentally altering tissue mechanics^11,12^. Our simulations reveal that lymph node expansion is governed by a mechanical operating window in which lymphocyte pressure, FRC contractility, and ECM elasticity must remain balanced to support robust and reversible tissue remodelling. Exceeding this operating window, either through excessive FRC contractility or ECM stiffening, progressively limits tissue expansion and alters the spatial distribution of FRC proliferation. Importantly, the spatial distribution of mechanical perturbations impacted lymph node remodelling to a greater degree than the total area of pathological alteration, highlighting the importance of balanced tissue mechanics in supporting lymph node expansion.

## Results

### Tissue morphometrics are maintained throughout lymph node expansion

The reticular network is often considered a single structure, but we now ask how cellular and non-cellular components cooperate and remodel during lymph node (LN) expansion. To investigate this, we immunised mice with incomplete Freund’s adjuvant/ovalbumin (IFA/OVA) and measured the reactive LN’s size, mass and stromal tissue morphometrics over the immunisation timecourse. We find that LNs increase in size and stay enlarged until day 14 (Figure 1a), with the mass increasing >3-fold (Figure 1b). The ECM conduit is ensheathed by the FRC network, physically separating it from the surrounding parenchyma (Figure 1c)^9,13,14^. We find that, at steady state, only 9% of the FRC network is not colocalised with Collagen I, largely due to a small subset of FRC branches lacking ECM, compared to 24% by day 5. FRC-ECM colocalisation is largely restored by day 9 (17%) and day 14 (13%), despite the LN remaining expanded (Figure 1d). We quantified areas between FRC branches in the LN paracortex (Figure 1e), the T cell-rich region of the LN, which increased significantly by 49.6% at day 5 (Figure 1f) and returned to steady state area by day 14, consistent with previous reports, attributing this to FRC network remodelling^2,3,15^. However, the eccentricity (deviation from circularity) of T cell zones was maintained throughout LN expansion, indicating uniform tissue expansion (Figure 1g,h). Previously, it has been reported that increased lymphocyte pressure in the LN causes the FRC network and associated ECM to stretch, causing some ECM structures to rupture^2,16,17^. Therefore, next, we characterised FRC and ECM thickness across LN expansion (Figure 1i-l). In agreement with previous reports, we find both cellular and non-cellular components become thinner at day 5 but gradually restore to steady state thickness by day 14. Collagen I thickness is significantly more variable at day 14 than at steady state (Extended Data Figure 1a). Together, these data show that the FRC and ECM networks do not spatially separate during tissue expansion. Tissue morphometrics such as T cell zone areas and eccentricities are restored and maintained independently of overall tissue size, raising the question of whether ECM dynamics are important for robust tissue remodelling.

**Figure 1.**
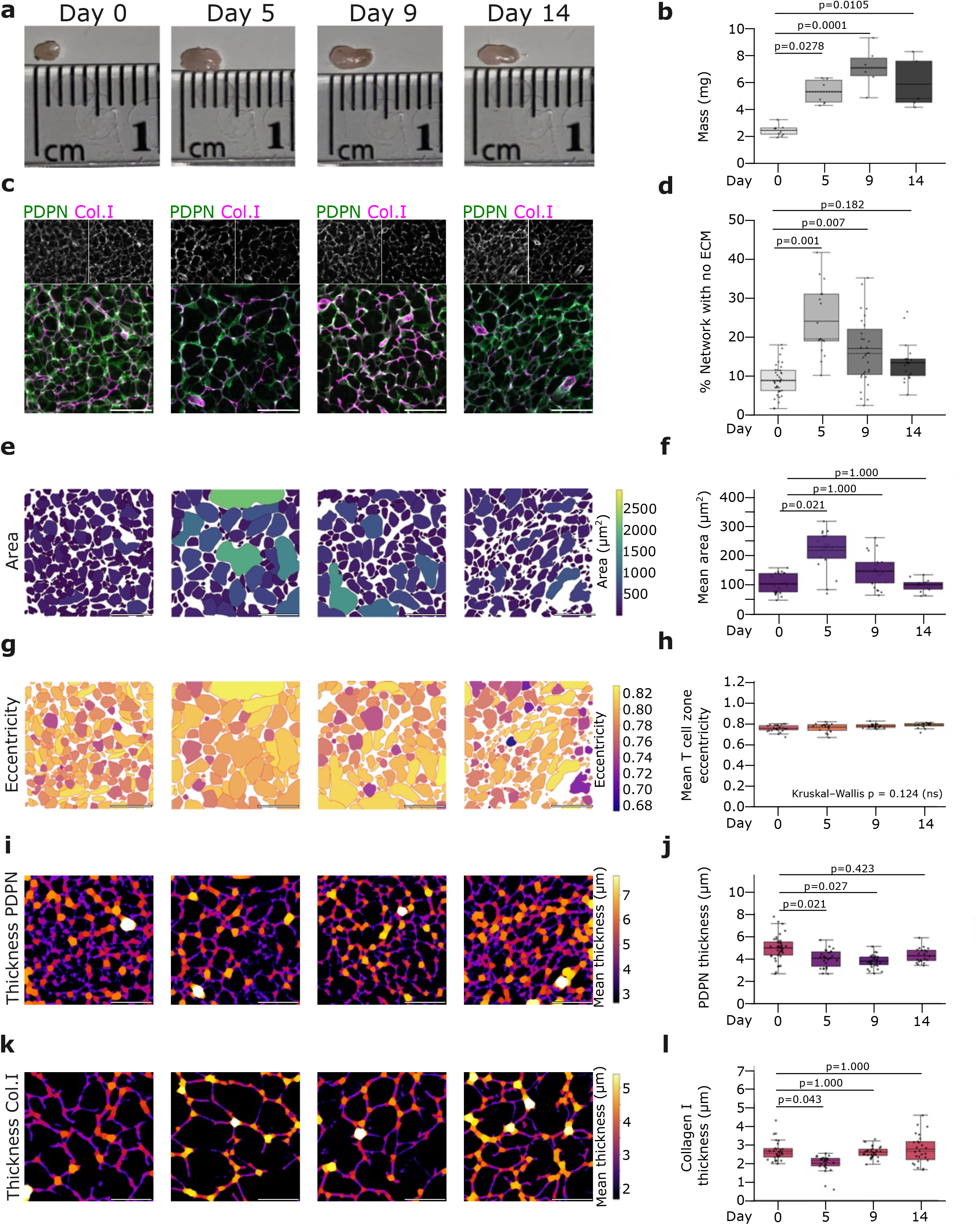
Tissue morphometrics are maintained throughout lymph node expansion. **a**, Representative images of LNs at day 0, 5, 9, 14 post IFA/OVA immunisation. **b**, LN mass (mg) after IFA/OVA immunisation. Each point represents one LN (N = 5-10 per timepoint). Box = IQR; whiskers = 1.5×IQR; solid line = mean; dashed line = median. Significance relative to day 0 was determined using Kruskal-Wallis test followed by Dunn’s multiple comparisons test; exact adjusted p-values are shown. **c**, Representative images of the FRC network in the LN during immunisation (PDPN, green and Collagen I, magenta). Scale bars, 50µm. **d**, Percent of FRC network not colocalised with ECM. Each point represents an individual ROI; statistics performed on LN-level means (n = 17-37 ROIs, N = 3-8 LNs per timepoint). Box = IQR; whiskers = 1.5×IQR; solid line = mean; dashed line = median. Significance relative to day 0 was determined using Kruskal-Wallis test followed by Dunn’s multiple comparisons test; exact adjusted p-values are shown. **e**, Napari segmentation of T cell zone areas during immunisation. Colour scale indicates area. Scale bars, 50µm. **f**, T cell zone area over immunisation timecourse. Each point represents an individual ROI; statistical analysis on LN-level means (n = 15-27 ROIs, N = 5-7 LNs per timepoint). Box = IQR; whiskers = 1.5×IQR; solid line = mean; dashed line = median. Box colour indicates median area (viridis scale). Significance relative to day 0 was determined using Kruskal-Wallis test followed by Dunn’s multiple comparisons test; exact adjusted p-values are shown. **g**, Eccentricity heat map of T cell zone areas during immunisation. Colour scale indicates eccentricity. Scale bars, 50µm. **h**, Mean T cell zone area eccentricity over immunisation timecourse. Each point represents an individual ROI; statistics on LN-level means (n = 15-27 ROIs, N = 5-7 LNs per timepoint). Box = IQR; whiskers = 1.5×IQR; solid line = mean; dashed line = median. Box colour indicates median eccentricity (plasma scale). Significance was determined using Kruskal-Wallis test; exact p-value shown. **i**, Heatmap of FRC network thickness (PDPN staining; colour scale, µm). Scale bars, 50µm. **j**, FRC network thickness (µm) over immunisation timecourse. Each point represents an individual ROI; statistics on LN-level means (n = 21-40 ROIs, N = 4-8 LNs per timepoint). Box = IQR; whiskers = 1.5×IQR; solid line = mean; dashed line = median. Box colour indicates median thickness (inferno scale). Significance relative to day 0 was determined using one-way ANOVA followed by Tukey’s multiple comparisons test; exact adjusted p-values are shown. **k**, Heatmap of Collagen I network thickness (colour scale, µm). Scale bars, 50µm. **l**, Collagen I network thickness (µm) over immunisation timecourse. Each point represents an individual ROI; statistics on LN-level means (n = 26-31 ROIs, N = 5-7 LNs per timepoint). Box = IQR; whiskers = 1.5×IQR; solid line = mean; dashed line = median. Box colour indicates median thickness (inferno scale). Significance relative to day 0 was determined using Kruskal-Wallis test followed by Dunn’s multiple comparisons test; exact adjusted p-values are shown.

### Extracellular matrix contributes to viscoelastic properties of the FRC network

It has been established that ECM stretches and ruptures during LN expansion^2,17^; however, what mechanical role ECM plays in LN architecture has not been tested. In a PKN2 knockout (KO) mouse model we report areas of ECM not colocalised with FRCs (Figure 2a). LNs in PKN2 KO mice expand more rapidly, suggesting that FRC-ECM mechanical association controls tissue expansion (Millward et al., https://doi.org/10.21203/rs.3.rs-4921177/v2). We find that in PKN2 KO LNs, Collagen I fibres are more curved at steady state (Figure 2b,c). After immunisation, Collagen I fibres become a straighter, consistent with stretching during LN expansion. To directly test the mechanical role of ECM in LN tissue mechanics, we enzymatically digested matrix components (Figure 2d). Collagenase is able to partially deplete conduit ECM, reducing Collagen I integrated density by 81% and second harmonic generation (SHG) maximum intensity by 35% (Figure 2e). Although ECM digestion does not significantly change the area or aspect ratio of a LN tissue cross section, we note a trend towards reduced area in cross-section (Figure 2f,g). To directly test whether ECM loss affects FRC network mechanics, we carried out laser ablations on the FRC network with and without ECM (Figure 2h). In control tissues, the recoil post laser ablation is restricted to areas directly adjacent to the ablated branch (Movies 1,2). In collagenase-treated network the recoil propagates further across the network, away from ablation site (Figure 2h). Likewise, the ablation site shows longer recoil times (brown bar) with greater total displacement distance (blue bar) (Figure 2i), in collagenase-treated tissues (Figure 2j). Further, recoil dynamics are more variable in collagenase-treated tissues (Figure 2k,l). This variability could arise due to uneven ECM digestion in the FRC network. Alternatively, intact ECM may homogenise the mechanical state of the FRC network and upon digestion, stronger and weaker network areas are exposed increasing recoil variation. From laser ablation displacement curves, we calculated initial recoil and elasticity-viscosity ratio of the reticular network (see Methods). The initial recoil is not significantly different between control and collagenase-treated tissues but tends towards a subtle increase (Figure 2m, control avg: 0.338 *µm*s^-1^; collagenase avg: 0.428 *µm*s^-1^), whereas the elasticity-viscosity ratio decreases by 28% upon collagenase treatment (Figure 2n). For the elasticity-viscosity ratio to decrease, viscosity must increase, elasticity must decrease, or both. We expect the main contributor to viscosity in the LN to be T cell packing density, which remains unaltered upon collagenase treatment (Extended Data Figure 2a,b). With viscosity unchanged, we therefore conclude that ECM contributes to FRC network elasticity at steady state.

**Figure 2.**
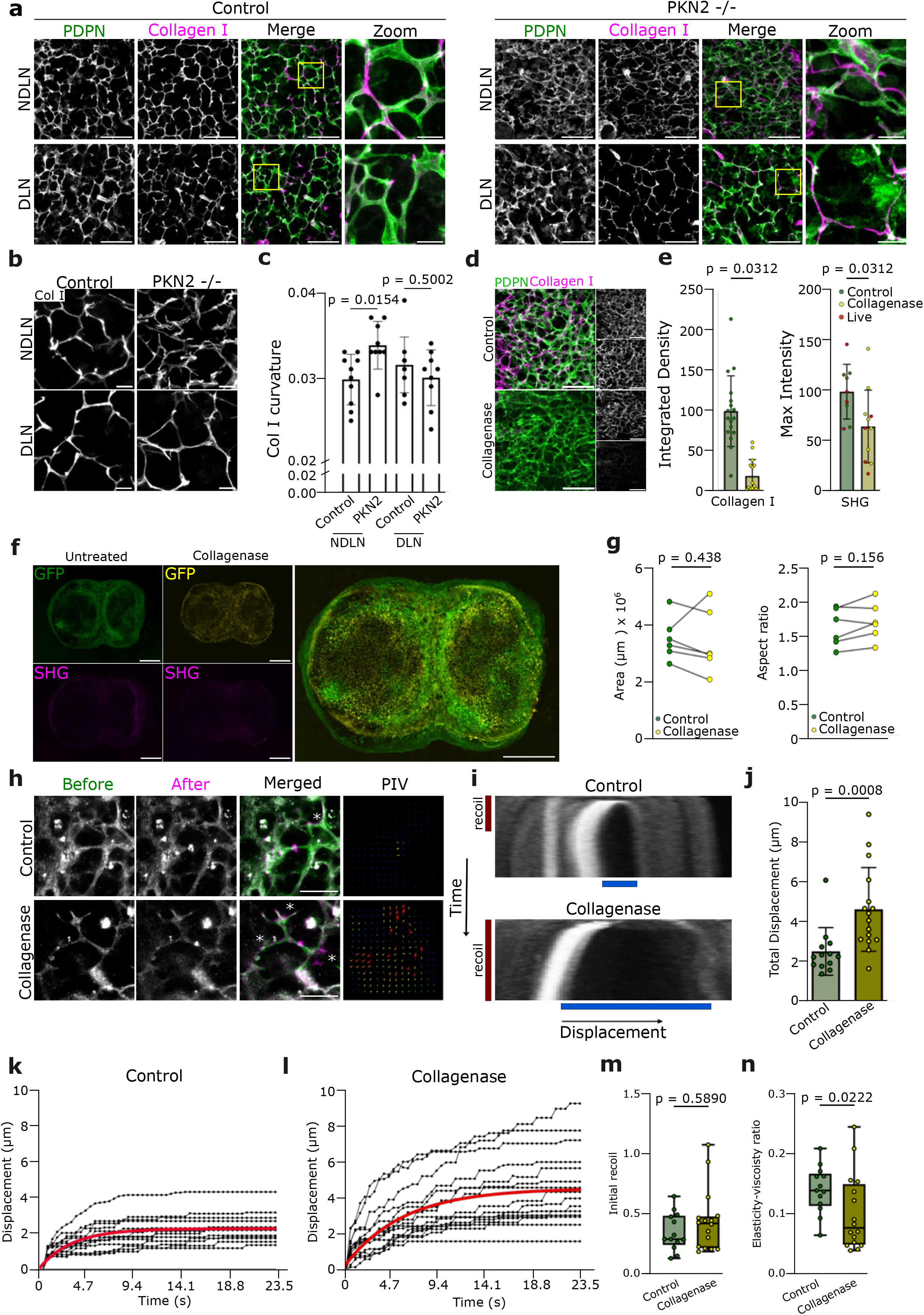
Extracellular matrix contributes to viscoelastic properties of the FRC network. **a**, Representative FRC network stained for PDPN (FRCs, green) and Collagen I (magenta) in control and immunised LNs from WT and PKN2 KO mice. Scale bars, 50µm; zoom-in, 10µm. **b**, Representative ECM network stained for Collagen I in WT and PKN2 KO mice. Scale bar, 10µm. **c**, Collagen I curvature quantified using TWOMBLI for WT control (**n**=10), WT immunised (**n**=10), PKN2 KO control (**n**=9) and PKN2 KO immunised (**n**=9) mice. **n** represents individual image ROIs, from N = 2 mice per genotype, each contributing one draining and one non-draining LN. Statistics: Ordinary two-way ANOVA followed by Šídák’s multiple comparisons test; adjusted exact p-values are shown. **d**, Representative control and collagenase-treated FRC networks, imaged using PDPN (FRC network, green) and Collagen I (magenta). Scale bar, 50µm. **e**, Integrated density of Collagen I (fixed tissue only; control n=16, collagenase-treated n=15 ROIs) and SHG signal (pooled fixed and live tissue; control n=9, collagenase-treated n=11 ROIs). Green and yellow dots = fixed-tissue control and collagenase-treated measurements, respectively; red dots = live-tissue SHG measurements. Statistics were performed on LN-level means (N = 5 paired LNs for each metric). Statistics: Exact one-tailed Wilcoxon matched-pairs signed-rank test; exact p-values shown. **f**, Representative tile scan (10x magnification) of live untreated (GFP, green) and collagenase-treated (GFP, yellow) LN tissue slice capturing SHG (ECM, magenta). Scale bar, 500µm. **g**, Quantification of LN tissue slice area and aspect ratio before (green) and after (yellow) collagenase treatment, showing no significant change (N = 5 independent LNs). Statistics: Exact two-sided Wilcoxon matched-pairs signed-rank test; exact p-values shown. **h**, Representative laser ablation of FRC networks in control and collagenase-treated samples, showing the network before ablation (green) and after ablation (magenta), with the corresponding PIV plot. Scale bar, 20µm. **i**, Representative kymographs showing network displacement (blue) following laser ablation in control and collagenase-treated samples, recoil time is shown in brown. **j**, Total displacement (µm) following laser ablations in control (green) and collagenase-treated samples (yellow). Statistics: Two-sided unpaired t-test with Welch’s correction; p-values shown. **k**,**l**, Recoil curves of network displacement for control (k) and collagenase-treated (l) samples. Red line indicates the mean. **m**,**n**, Initial recoil and Elasticity-viscosity ratio respectively in control (green) and collagenase-treated (yellow) samples. Statistics: Exact two-sided Mann-Whitney U test; exact p-values shown. For panels **j**,**k**,**l**,**m**,**n**, each data point represents an individual laser ablation (n=13 control, n=16 collagenase) from N = 4 independent experiments.

### A 2D vertex model of FRC network captures the mechanical interplay between FRCs, T cells, and ECM in the lymph node

FRCs exist as an intrinsically contractile cellular network, surrounded by T cells which create pressure in the LN and promote organ growth^2,3^. These two opposite forces are balanced at steady state, and their magnitudes determine the overall organ size^2,3^. Since in 3D space, T cells move freely between connected network structures (Figure 3a); we conceptualise T cell pressure as uniform across all T cell areas. ECM’s contribution to LN size has not been addressed experimentally; however, our *ex vivo* experiments show that ECM significantly affects FRC network mechanics (Figure 2h-n). Here, we adapt a 2D vertex model^18^ that captures the FRC network mechanics, T cell pressure, and ECM mechanics (Figure 3b). In our model, vertices represent FRC branch points, edges represent individual FRC branches, and polygons represent T cell areas (Figure 3b). The polygons exert forces to reach their preferred area, and the edges are contractile, reflecting the LN force balance. To assess how ECM and FRCs contribute to LN mechanics, we compared two energy equations. Both capture the pressure arising from T cell density with an area elasticity term, where each T cell zone has a preferred area, 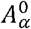 (red), toward which it tends. We then modelled the mechanical interaction of ECM and FRCs in two different ways (Figure 3, Models 1 and 2). In Model 1, length elasticity term, *K*, is split into the cellular component, 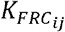, and the ECM contribution, 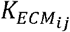 which can have non-equal values. We assume in Model 1 that each edge has a preferred length, 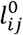, which emerges from combined mechanical contribution of ECM and FRCs. In Model 2, we separate the cellular and ECM mechanical contributions into two independent energy terms. Since regions of the ECM can maintain structural integrity independently of the FRC network in PKN2 KO mice (Figure 2a), the cellular and extracellular components may be captured best by distinct mechanical terms in the model. In Model 2, length elasticity term, 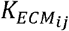, with an associated preferred length reflects ECM mechanics. FRCs are modelled with a line tension term, 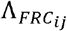, which reflects FRC actomyosin contractility. The observed length of an FRC branch in Model 2 therefore emerges from the balance between cellular contractility, ECM’s preferred length and stiffness, external pressure, and connectivity within the reticular network. Figure 3c illustrates how the forces in Models 1 and 2 act on the FRC network under compressed, relaxed, and stretched conditions. In Model 1, edges exert forces to return to their preferred length in compressed and stretched scenarios. In relaxed scenario, *l*_*ij*_ *= l*_*0*_, so length elasticity term becomes zero. In Model 2, the line tension term, capturing FRC contractility, is not dependent on preferred length and is therefore always positive, acting to minimise the edge length in all scenarios. This is akin to a contractile cell minimising its surface area by adopting a spherical shape.

**Figure 3.**
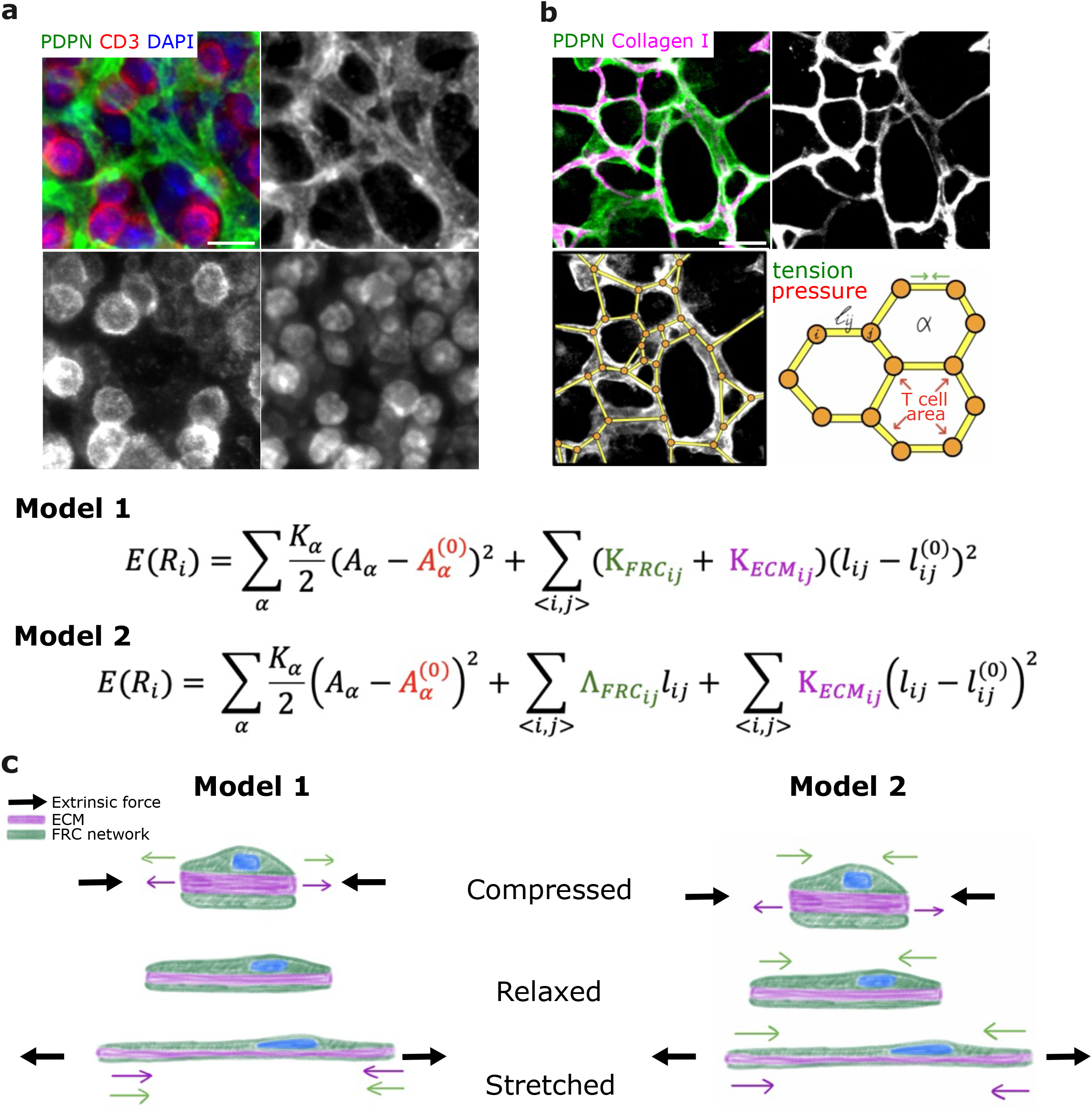
Modelling independent mechanical contributions of FRCs and ECM better recapitulates FRC network mechanics. **a**, Representative FRC network stained for PDPN (FRC network, green), CD3 (T lymphocytes, red) and DAPI (nuclei, blue). Scale bar, 20µm. **b**, FRC network stained for PDPN (FRCs, green) and Collagen I (magenta). Scale bar, 20µm. A 2D vertex model is overlayed on the FRC network, with edges aligned to FRC branches and vertices aligned to network branch points. Schematic showing simplification of the FRC network into a 2D vertex model, incorporating tension from FRCs and pressure from T cells. Model 1 treats FRC and ECM contributions as a combined effective edge stiffness, whereas Model 2 separates FRC contractility from ECM elasticity (see Methods). **c**, Schematic illustrating the mechanical response of Model 1 and Model 2 under compression, relaxation and stretch. In Model 1, FRC and ECM contributions are combined into a single effective elastic term, whereas in Model 2, FRC contractility and ECM elasticity are represented as separate contributions. Black arrows indicate extrinsic force.

### FRC contractility and ECM elasticity make distinct contributions to reticular network mechanics

To evaluate which model better captures LN mechanics, we parameterised area elasticity, length elasticity and viscosity for both Models, and line tension for Model 2. Each parameterisation was benchmarked by comparing initial recoil and elasticity-viscosity ratio between *in silico* and *ex vivo* laser ablations (Figure 4a). Model 2 provided a closer overall fit to *ex vivo* data than Model 1, achieving a lower minimum Mahalanobis distance of 0.92 vs 1.49 and a greater number of parameter combinations within a Mahalanobis distance of 1.6 (Figure 4b). We next tested whether both models accurately capture the FRC network mechanics observed following ECM loss. For Model 1, where the length elasticity term is split between 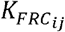 and 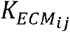, we conducted *in silico* laser ablations across a range of 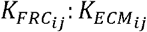 ratios with 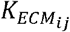set to 0 to mimic ECM loss. A 1:5 ratio most closely recapitulated *ex vivo* collagenase laser ablation data and thus was chosen for further analysis (Extended Data Figure 3, Figure 4c). *In silico* laser ablations show greater variation in Model 2 than Model 1 (Figure 4c,d), which more closely matches *ex vivo* data. *Ex vivo*, following ECM loss, initial recoil does not significantly change, the elasticity-viscosity ratio decreases, and total displacement increases. *In silico*, initial recoil decreases in both models upon collagenase simulation, but the decrease is less in Model 2 than in Model 1 (Figure 4e). The elasticity-viscosity ratio decreases in both models, consistent with *ex vivo* data (Figure 4f). Total vertex displacement decreases in Model 1 upon collagenase simulation but increases in Model 2, consistent with *ex vivo* data (Figure 4g). Model 1 appears homogeneous when initialised, whereas Model 2 shows T cell area variability despite both models using uniform parameters (Figure 4i,j, Control). This is echoed in the force plots, where Model 2 shows higher force magnitudes than Model 1 (Figure 4i,j, Force plot). Upon collagenase treatment *ex vivo*, we observe lengthening of FRC branches and enlargement of T cell areas (Figure 4h), a phenotype captured by Model 2 but not Model 1 (Figure 4i,j, Collagenase). These findings indicate that Model 2 more accurately captures FRC network mechanics than Model 1, and suggest that the reticular network should not be thought of as a single mechanical entity; instead, FRCs and ECM contribute distinct mechanical functions, with active cellular contractility coupled to an elastic extracellular scaffold. Model 2 is therefore used for the remainder of the study.

**Figure 4.**
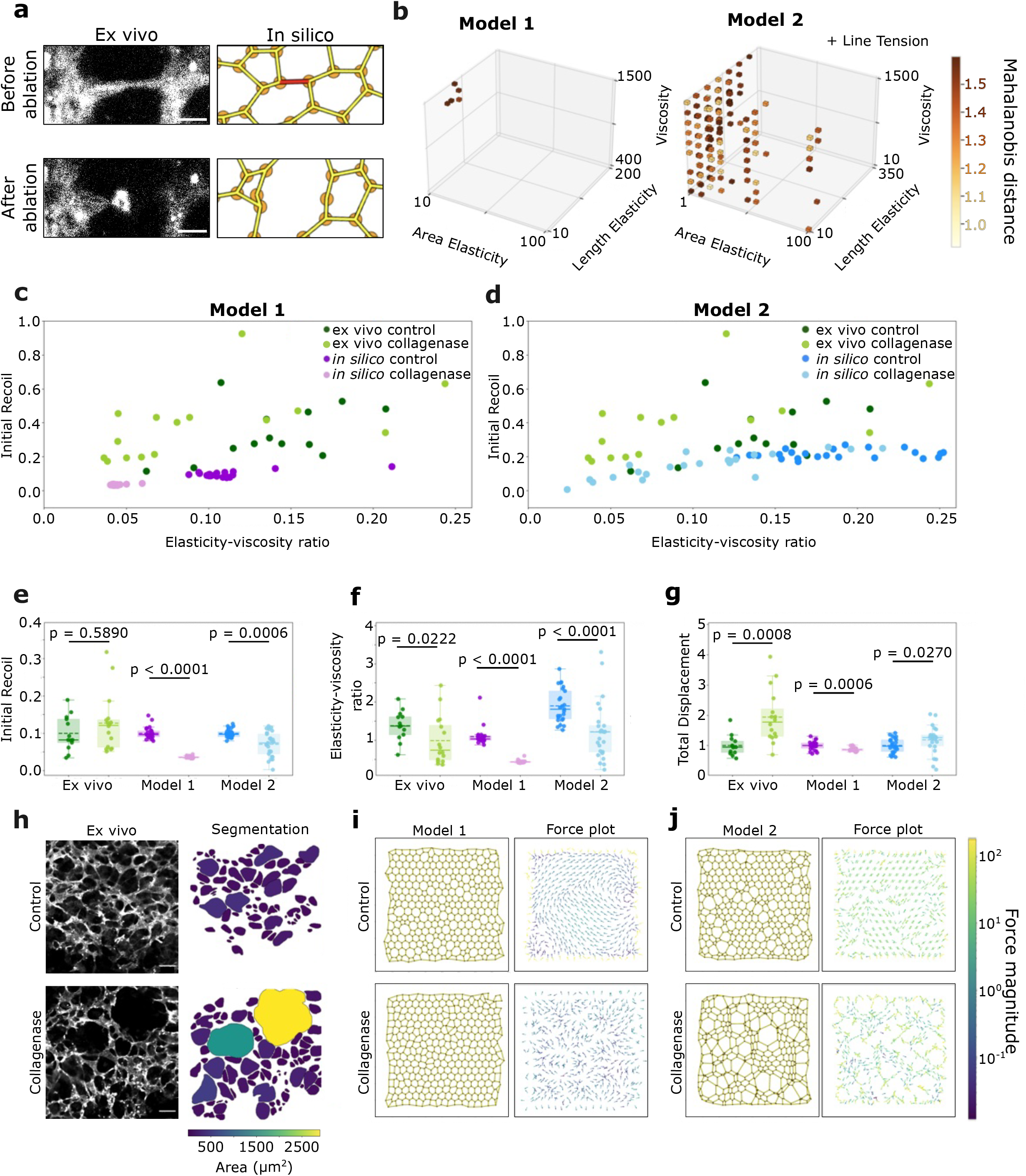
Modelling independent mechanical contributions of FRCs and ECM better captures FRC network mechanics. **a**, Representative *ex vivo* and *in silico* FRC networks shown before and after ablation. Scale bar, 5µm. **b**, Three-dimensional projection of the parameter space for Model 1 and Model 2. Only parameter combinations with a Mahalanobis distance < 1.6 are shown. Points are positioned according to area elasticity, length elasticity, and viscosity and coloured by Mahalanobis distance. In Model 2, line tension was additionally varied during the parameter sweep but is omitted from this projection. **c**, Scatter plot of initial recoil against elasticity-viscosity ratio for *ex vivo* laser ablation of control FRC networks (dark green) and collagenase-treated FRC networks (light green), and for *in silico* laser ablation of control models (dark magenta) and collagenase-simulated models (light magenta). Each point represents a single ablation. *Ex vivo*: control **n**=13 and collagenase-treated **n**=16 from N = 4 independent experiments; *in silico*: control n=25 and collagenase-simulated n=25 from N = 5 independent simulations. **d**, Scatter plot of initial recoil against elasticity-viscosity ratio for *ex vivo* and *in silico* laser ablations, for Model 2 under control (dark blue) and collagenase-simulated (light blue) conditions; **n** values are as in **c. e**,**f**,**g**, Comparison of *ex vivo* data (green) with Model 1 (magenta) and Model 2 (blue) under control (dark) and collagenase conditions (light). **e**, Initial recoil. **f**, Elasticity-viscosity ratio. **g**, Total network displacement following laser ablation. Each dot represents a single laser ablation. *Ex vivo* data comprise n = 13 control and n = 16 collagenase-treated ablations from N = 4 independent experiments. *In silico* data comprise n = 25 control and n = 25 collagenase-treated ablations from N = 5 independent simulations. Statistics: depending on normality and variance, an exact two-sided Mann-Whitney U test or two-sided unpaired t-test with Welch’s correction was used (see Methods); p-values shown. **h**, Representative images of control and collagenase-treated FRC networks. Scale bar, 20µm. Napari segmentation of T cell areas is shown for comparison (colour represents area). **i**,**j**, Representative Models 1 and 2 configurations under control and collagenase-simulated conditions, shown alongside the corresponding force plots.

### Modelling predicts that mechanical properties of ECM regulate lymph node tissue expansion

A central question in LN expansion is how the FRC network adapts to it while maintaining structural integrity. Multiple factors contribute, including FRC division and apoptosis. To incorporate division and apoptosis into our model, we tested different mechanisms, comparing the resulting network geometry and topology to *in vivo* FRC networks (Extended Data Figures 4,5). Division by splitting the vertex with the largest summed edge length, combined with apoptosis by random edge collapse, most closely recapitulates FRC network architecture (Extended Data Figure 6).

Having introduced FRC division and apoptosis into the model, we next systematically tested the mechanical contributions of pressure, tension, cell division, and ECM mechanics to tissue expansion. We modelled the increase in immune cell numbers as increased area elasticity, *K*_*a*_, and increased preferred area, *A*_*0*_. To model FRC network relaxation and stretching, we decreased line tension, 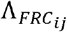. As LN expands, FRCs stretch, which triggers their proliferation^3^, an emergent property in the model (Figure 5a). To permit tissue expansion, we decreased outer vertex viscosity, modelling the previously reported softening of the LN capsule^2^. We then systematically tested each parameter *in silico* to quantify their effect on overall tissue expansion and architecture (Figure 5b-d). We find that increased pressure alone is insufficient to drive physiological tissue expansion (Figure 5b,c; Extended Data Figure 7, Movie 3). However, increased pressure together with reduced network tension permits growth of the simulated tissue and accurately replicates T cell area fold change *ex vivo* from Day 0 to Day 5 of the immune response (Figure 5d). No significant additional growth is achieved by allowing longest edges to instruct FRC division (Figure 5c,d). We compared simulated expansion with Day 9 *ex vivo* data, when FRC division occurs, division alone was insufficient to reproduce the observed reduction in individual T cell areas (Figure 5d). This may reflect model’s exclusion of cellular processes such as FRC branching (Millward et al., https://doi.org/10.21203/rs.3.rs-4921177/v2). *In vivo*, FRC network is predicted to relax globally at the start of an immune response^19,20^. We used our model to test whether the network requires uniform tension reduction, randomly assigning a 0.5-1.0 proportion of edges to remain at higher tension as pressure increased (Figure 5e). Fixing as few as 10% of edges at higher tension fully constrains tissue growth, whereas a network with 99% relaxed edges expands substantially, though less than a fully relaxed network (Figure 5f, Movie 4). Complete FRC network relaxation (99-100%) promoted greater cumulative FRC division than partial relaxation (Figure 5g). Thus far, these simulations maintain homeostatic ECM stiffness, predicting that tissue expansion can occur without disrupting the underlying ECM. Our experimental and *in silico* data predict that the ECM resists cellular contractile forces and/or lymphocyte pressure during tissue expansion (Figures 2-4), providing a mechanism to control expansion and preserve tissue structure. To assess the role of ECM mechanics in LN expansion, we modelled progressive ECM softening or stiffening. ECM softening promoted tissue expansion, whereas ECM stiffening increasingly constrained tissue growth and reduced FRC division (Figure 5i,j, Movie 5). Supplementary Table 3 summarises parameters used in Figure 5b-j. These findings predict that the mechanical properties of ECM are important for LN remodelling and that disease states involving ECM loss or fibrotic stiffening would severely impair tissue remodelling and expansion (Figure 5i), and limit FRC division (Figure 5j).

**Figure 5.**
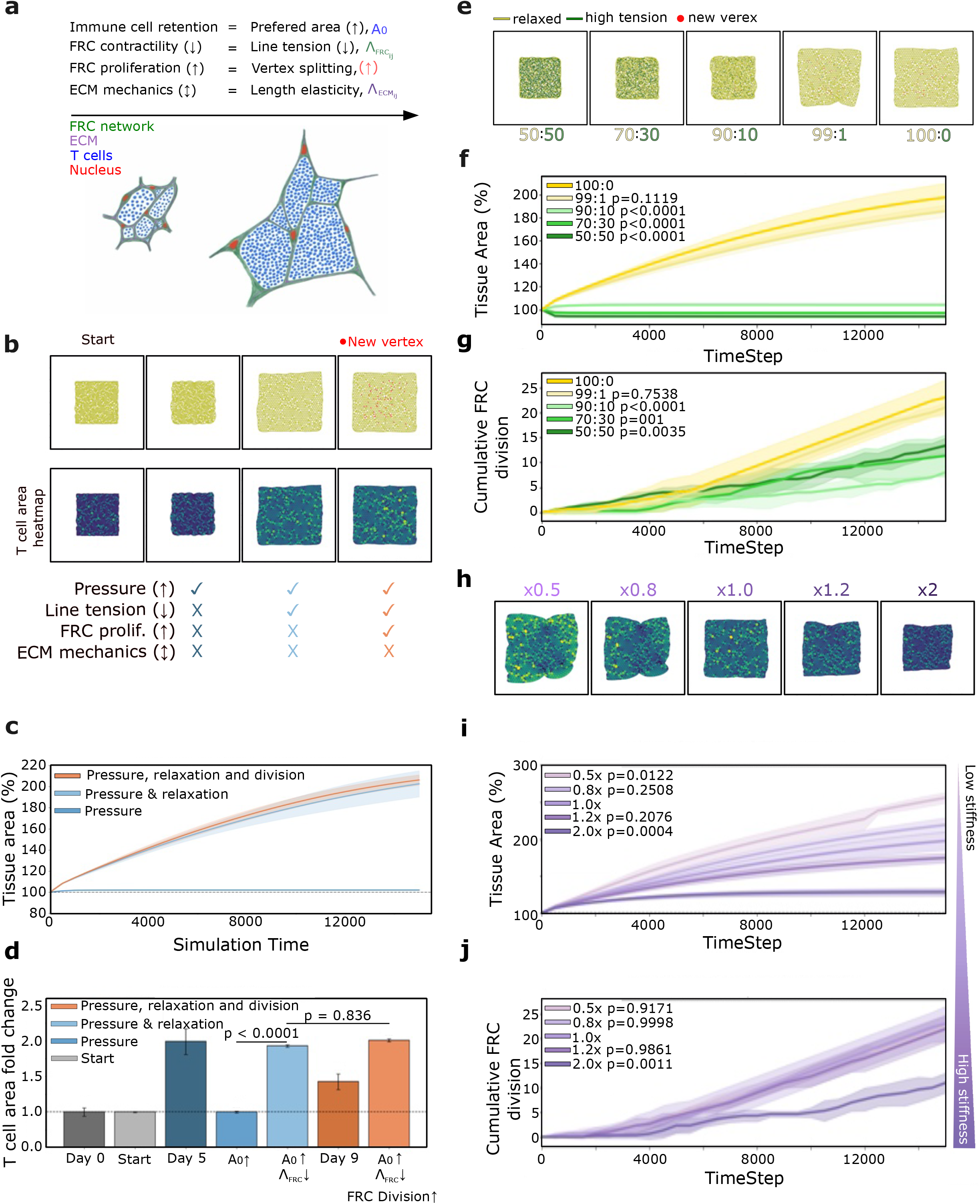
Modelling predicts that mechanical properties of ECM regulate lymph node tissue expansion. **a**, Schematic illustrating the mechanical representation of biological processes during LN expansion. Immune cell retention = increased preferred area (*A*_*0*_, blue); reduced FRC contractility = decreased line tension 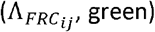; FRC proliferation = an emergent property of the model (red); ECM mechanics = changes in length elasticity (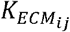, purple). **b**, Representative expansion simulations showing the initial network (Start), increased pressure alone (*A*_*0*_ x8, *K*_*a*_ = 50), increased pressure with reduced FRC tension (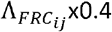, *A*_*0*_ x8, *K*_*a*_=25), and increased pressure with reduced FRC tension and FRC proliferation. Corresponding T cell area heatmaps are shown below each condition. **c**, Tissue area (%) over time for each expansion condition. **d**, Endpoint T cell area fold change for each expansion condition. *Ex vivo* Day 0 and the initial simulation state were each normalised to 1. Pressure-driven expansion with or without FRC relaxation is compared with Day 5, whereas pressure-driven expansion with FRC relaxation and proliferation is compared with Day 9 (*ex vivo* data n=13-23 ROIs, N=3-6 LNs). Bars show mean ± SEM. Statistics were performed on *in silico* groups only (ordinary one-way ANOVA followed by Tukey’s multiple-comparisons test); exact p-values shown. *Ex vivo* data shown for reference only. **e**, Endpoint network configurations with increasing proportions of relaxed FRC edges (yellow) to high-tension edges (green) (50:50, 70:30, 90:10, 99:1 and 100:0). **f**,**g**, Tissue area increase (%) and cumulative FRC division, respectively, over time for each proportion of relaxed FRC edges. **h**, Representative endpoint T cell area heatmaps for simulations with increasing ECM stiffness. **i**,**j**, Tissue area increase (%) and cumulative FRC division, respectively, over time for simulations with varying ECM stiffness. For all time-course plots, lines indicate the mean and shaded regions indicate s.d. Unless otherwise stated, each simulation condition reflects N = 3 independently initialised simulations. **f**,**g**,**i**,**j**, Statistics: Ordinary one-way ANOVA followed by Dunnett’s multiple-comparisons test comparing each condition with the corresponding control; exact p-values shown.

### The spatial organisation of mechanical pathology determines lymph node robustness

Tumour-draining lymph nodes (TDLNs) are sites of early metastatic spread and are used clinically as a prognostic indicator of tumour progression^21^. TDLNs undergo phenotypic and immunological changes prior to metastatic seeding, but the extent and functional significance of these alterations remain incompletely understood. In LNs from breast cancer patients, we observe geometric changes in the FRC network architecture, which predict patient outcome^22^ (Figure 6a). Chronically inflamed murine TDLNs also exhibit stromal phenotypic changes accompanied by FRC network disruption and progressive aberrant ECM deposition^12^. We find that the FRC network in TDLNs exhibits patches of increased α-smooth muscle actin (αSMA) expression and decreased PDPN prior to metastatic seeding (Figure 6b). αSMA is a functional marker of contractile myofibroblasts ^23^, which are known to deposit ECM with altered, stiffer properties^24^. Because these processes occur concurrently *in vivo*, it is experimentally difficult to isolate their individual mechanical contributions or the effects of their magnitude and spatial distribution. We therefore used our model to independently vary FRC hypercontractility 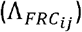 or ECM stiffness 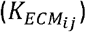 across different proportions (10, 30 and 50%) and spatial distributions (1, 10 and 50 patches), to predict their effects on LN expansion and network architecture. Despite altering the same proportion of the FRC network, perturbation distribution markedly influenced how deformation propagated through the tissue (Figure 6d, Movie 6). Both the extent and magnitude of mechanical perturbations progressively limited LN expansion for both FRC hypercontractility and ECM stiffening (Figure 6e-h). At higher FRC contractility (x1.5), and both ECM stiffening perturbations (x3 and x5), dispersed perturbations (10, 50 patches) impaired expansion more than localised perturbations (1 patch) when the same proportion of the network was affected (Figure 6f-h). However, at lower levels of FRC hypercontractility (×1.2), no significant differences were observed between different patch distributions (Figure 6e). These findings predict that localised mechanical perturbations are buffered by surrounding tissue, but the buffering is lost as pathology becomes spatially dispersed. For example, under five-fold ECM stiffening, perturbing 10% of the network across 10 dispersed patches impaired expansion to a similar extent as perturbing 50% within a single patch (Figure 6h). Hypercontractility and increased ECM stiffness produced distinct patterns of tissue impairment, with ECM stiffening exerting a stronger mechanical effect than comparable increases in contractility (Extended Data Figure 8a). Although mechanical perturbations consistently impaired expansion, their effects on global tissue shape were more variable and did not scale straightforwardly with perturbation magnitude or patchiness (Extended Data Figure 8b). Cumulative divisions broadly increased with final tissue area (Figure 6i-l), but some simulations reaching a similar final size accumulated markedly different numbers of divisions, indicating that mechanical perturbations alter division patterns independently of overall tissue expansion (Figure 6e-l, Extended Data Figure 8c). The majority of divisions occurred outside the perturbed regions (Figure 6m-p, Extended Data Figure 8d), indicating that mechanical defects propagate their impacts beyond the local perturbed tissue (Figure 6m-p).

**Figure 6.**
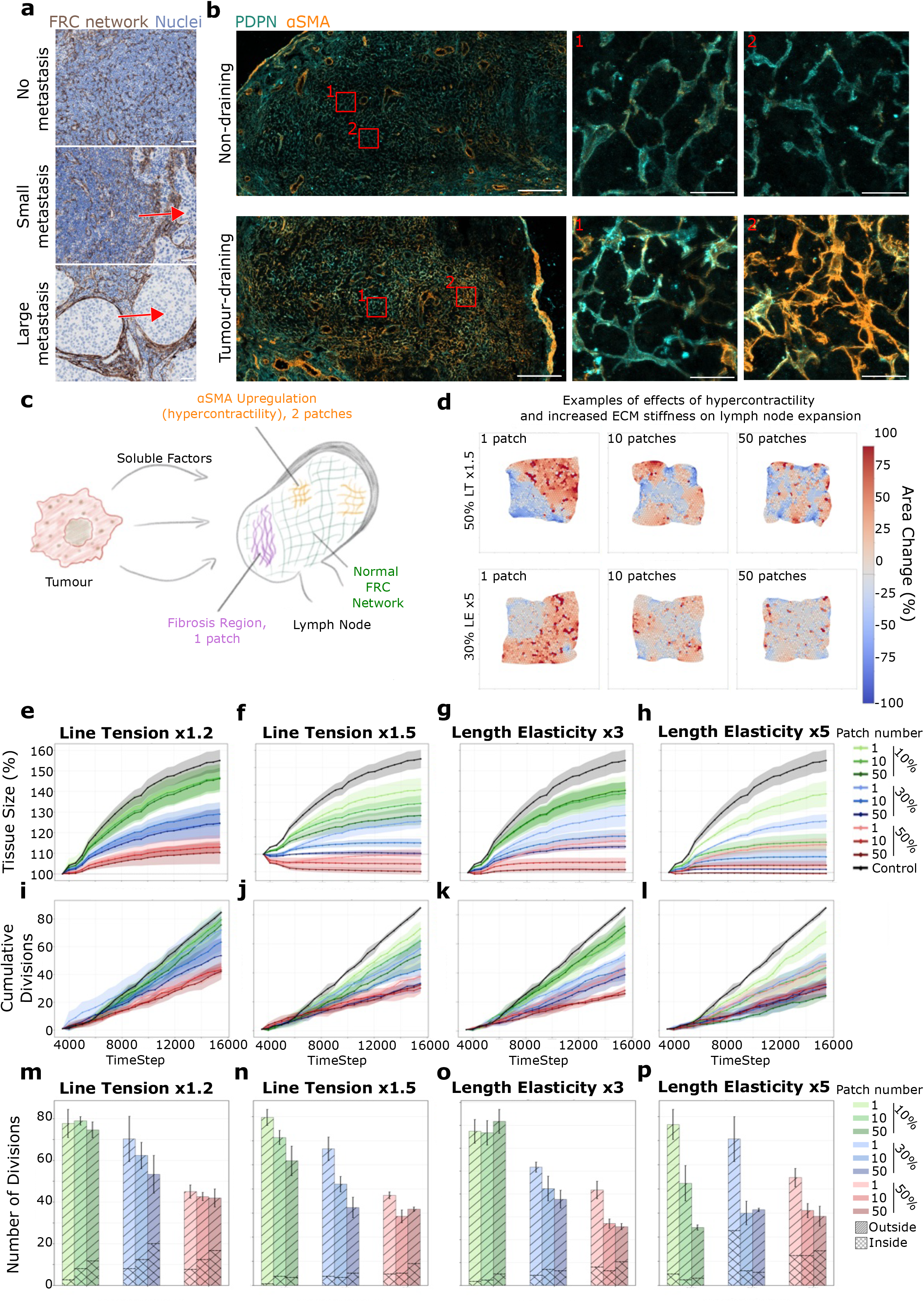
The spatial organisation of mechanical pathology determines lymph node robustness. **a**, Representative human LN sections with no metastasis, small metastasis and large metastasis, stained for PDGFRβ (brown) and nuclei (blue). Red arrow = metastatic deposit. Scale bars, 50µm. **b**, Representative tile scans of murine non-draining and tumour-draining LNs (TDLNs) stained for PDPN (cyan) and αSMA (orange). Scale bars, tile scan, 200µm; zoom-in, 20µm. **c**, A schematic showing effects of tumour soluble factors on TDLN, including ECM fibrosis and FRC hypercontractility. **d**, Representative heatmaps of local T cell area change during tissue expansion for simulations with increasing magnitudes and spatial distributions of FRC hypercontractility (line tension (LT) ×1.5) and ECM fibrosis (length elasticity (LE) ×5). Simulations were initiated from the N = 3 expansion checkpoints used in Figure 5 (simulation step 3,500). **e**-**l**, Tissue size (%) (**e**-**h**) and cumulative FRC divisions (**i**-**l**) over time for simulations with increasing magnitudes and spatial distributions of FRC hypercontractility (LT ×1.2 and ×1.5) or ECM fibrosis (LE ×3 and ×5). Simulations were performed with 10%, 30% or 50% of the network mechanically perturbed, distributed across 1, 10 or 50 patches, as indicated. Black lines = control simulations, no mechanical alterations. Lines = mean, shaded regions = s.d. **m**-**p**, Total number of FRC divisions occurring inside and outside mechanically perturbed regions in simulations with increasing magnitudes and spatial distributions of FRC hypercontractility (**m**, LT ×1.2; **n**, LT ×1.5) or ECM fibrosis (**o**, LE ×3; **p**, LE ×5). Error bars indicate s.d. Statistics: For comparisons of patch number (1, 10 or 50 patches) within each perturbation magnitude, ordinary one-way ANOVA followed by Tukey’s multiple-comparisons test was used. Comparisons between control and pooled perturbation magnitudes (10%, 30% and 50%) were performed using ordinary one-way ANOVA followed by Tukey’s multiple-comparisons test; p-values shown in Supplementary Table 2. Each perturbation condition in **e-p** reflects N = 3 simulations, using the same expansion checkpoints as in **d**.

## Discussion

Here, we investigate the mechanical determinants of lymph node homeostasis and expansion and develop an *in silico* biomechanical model of the lymph node, integrating the mechanical forces arising from immune cells, FRCs, and conduit ECM. This framework enables individual mechanical roles of each component to be systematically and locally dissected while remaining grounded in experimental measurements. Previous work has primarily focused on the biology of FRCs and their adaptation during immune responses, with the reticular network typically treated as a single mechanical entity. Our *ex vivo* tissue perturbations demonstrate that the conduit ECM provides the dominant elastic force in the lymph node, buffering FRC-generated contractility. Our work thus separates the mechanical functions of FRCs and the ECM, demonstrating that tissue expansion emerges from interactions between active cellular contractility and an elastic extracellular scaffold. Consistent with these observations, only computational models that assign distinct mechanical roles to FRCs and the ECM reproduce the recoil dynamics observed experimentally following laser ablation. These findings demonstrate that the reticular network should not be viewed as a single mechanical entity, but rather as two distinct yet mechanically coupled components whose interactions determine tissue-scale behaviour. More broadly, this framework establishes the lymph node as a uniquely tractable system for investigating how active cellular forces and passive extracellular mechanics cooperate to produce rapid, cyclical tissue remodelling in adult organs.

Our simulations further reveal that successful lymph node expansion depends on tissue-scale mechanical integration across the reticular network. Local increases in lymphocyte pressure, for instance around high endothelial venules (HEVs), and local reductions in FRC contractility following CLEC-2/PDPN signalling are likely to initiate lymph node remodelling. However, local perturbations alone are insufficient to drive homogeneous organ expansion. Instead, successful remodelling requires coordinated relaxation throughout the reticular network, allowing local mechanical perturbations to be integrated into a coherent tissue-wide response. How this transition from local biochemical signalling to tissue-wide mechanical remodelling is achieved remains an important unanswered question. One possibility is that local reduction in FRC contractility alters tension within the conduit ECM, enabling changes in force to propagate throughout the reticular network. Such force transmission could provide neighbouring stromal cells with mechanical cues to modulate their own contractility, thereby amplifying local immune-driven signals into coordinated organ-scale remodelling. This hypothesis provides a potential mechanism by which local immune activation in the lymph node could be translated into coordinated tissue-scale remodelling across the lymph node.

Consistent with this concept, local increases in FRC contractility or ECM stiffening disrupted tissue architecture and FRC proliferation beyond the perturbed regions, demonstrating that local mechanical changes propagate throughout the reticular network. This raises the possibility that pathological stromal remodelling changes lymph node architecture, not only through local alterations in matrix composition and cellular contractility, but also by redistributing normal patterns of FRC turnover. The importance of reticular network architecture for immune cell migration, chemokine organisation, and conduit-mediated antigen transport is well established. Our findings provide a mechanical framework for understanding how this architecture is robustly preserved during the dramatic yet reversible expansion of the lymph node and, conversely, how pathological alterations in tissue mechanics may ultimately compromise immune function. Thus, this computational framework provides a powerful platform for investigating how mechanics shapes lymph node organisation and function during homeostasis, immune responses, and disease.

## Methods

### Mice

All animal experiments were performed in accordance with national and institutional guidelines for animal welfare and were approved by the relevant institutional ethics committee and the UK Home Office. Mice were maintained under specific pathogen-free conditions. Wild-type C57BL/6J mice were purchased from Charles River Laboratories. Pdgfrα-mGFP-CreERT2 mice^3,25^, in which membrane-targeted GFP is expressed under the control of the endogenous Pdgfrα promoter, were used to visualise PDGFRα-expressing stromal cells. PKN2^fl/fl^ x Rosa26^CreERT2/WT^ x Rosa26^mT/mG^ mice (Rosa26^ind.ΔPkn2^) and PKN2^WT/WT^ x Rosa26^CreERT2/WT^ x Rosa26^mT/mG^ littermate controls (cre-only controls), in which tamoxifen-induced Cre recombination disrupts PKN2 function in Cre-expressing cells, were gifted by Dr Angus Cameron, Queen Mary University of London, and have been described previously^26^ (Millward et al., Research Square). Female and male mice aged 8-15 weeks were used for experiments unless otherwise stated. Animals were randomly assigned to experimental groups. Cre-mediated recombination in Pdgfrα-mGFP-CreERT2 and Rosa26^ind.ΔPkn2^ mice was induced by intraperitoneal injection of tamoxifen (reconstituted at 22 mg/mL in corn oil) at a dose of 2 mg per 20g body weight, administered using a 27-gauge needle for 4 consecutive days. The injection side was alternated between the left and right intraperitoneal cavity each day to minimise local irritation, and mice were weighed before and after tamoxifen administration (Millward et al., Research Square).

### Immunisation protocol

100µl of an emulsion of Ovalbumin (OVA) and Incomplete Freund’s Adjuvant (IFA) (Hooke Laboratories) was injected subcutaneously in the right flank of every mouse, between ribs and hip (100µg OVA per mouse).

### Lymph node isolation

Carbon dioxide (CO_2_) chamber was used to euthanise all mice, set to 5 minutes of CO_2_ exposure, followed by 1 minute of CO_2_ absence. Cervical dislocation was used as confirmation of death. Mice were euthanised at desired timepoints after immunisation with IFA/OVA (day 0, day 5, day 9, day 14). Inguinal and axillary lymph nodes were identified and any fat covering them was removed. Then lymph nodes were gently lifted with forceps from below, avoiding compression damage. Lymph nodes were either placed directly into PBS on ice for fixation or into complete RPMI 1640 medium (ThermoFisher Scientific) for live vibratome sectioning. The complete medium consisted of RPMI 1640 supplemented with 10% fetal bovine serum (FBS; Sigma-Aldrich), 1% Insulin-Transferrin-Selenium (ITS; ThermoFisher Scientific), and 1% Penicillin-Streptomycin (10,000U/mL; ThermoFisher Scientific). Prior to further processing, lymph nodes were weighed using a Sartorius Quintix™ Semi-Micro Balance (Fisher Scientific).

### Vibratome sectioning of lymph nodes

Vibratome sectioning was optimised in the laboratory, building upon previous methods^3^. Lymph nodes were removed from PBS/medium and dabbed onto a tissue to remove any excess moisture. LNs were placed at the bottom of a mould and UltraPure™ low melting point agarose (Thermo Fisher Scientific), 3% w/v, kept at 37°C, was poured on top (to investigate the effect of collagenase on LN tissue shape, the LN slice was instead embedded in 1% agarose to allow the tissue to change shape). The moulds were placed for a few minutes on ice for the agarose to set. The vibratome chamber was filled with ice cold PBS and ice was placed in the surrounding area to keep PBS cold during sectioning. Once set, LN blocks were extracted from the mould and excess agarose around the LN was removed with a scalpel. This is done to minimise distance over which the vibratome needs to cut and therefore decreases sectioning time. The LN block was then superglued onto the removable vibratome cutting stage. The stage was fitted back into the cutting chamber using the magnetic adherence. Sections were cut using a Leica VT1200S vibratome with settings of 1.5mm amplitude, 0.3mm/s speed, and a section thickness of 200µm. A new blade (Avantor) was fitted into the blade holder for each session and lowered until the white alignment line on the rotating blade mount became visible. The first and last tissue slices were excluded, as they predominantly consisted of the lymph node capsule. The central 2-4 slices, depending on the size of the lymph node, were collected in complete medium comprising RPMI 1640 (ThermoFisher Scientific) supplemented with 10% fetal bovine serum (FBS; Sigma-Aldrich), 1% Insulin-Transferrin-Selenium (ITS; ThermoFisher Scientific), and 1% Penicillin-Streptomycin (10,000 U/mL; ThermoFisher Scientific). Slices were immediately imaged using a microscope equipped with a chamber maintained at 37°C and 10% CO_2_.

### ECM digestion

10mg of collagenase type II (Worthington, LS004176) was weighed and dissolved in 1mL of complete medium to prepare a 10mg/mL stock solution. Following vibratome sectioning, lymph node slices were either transferred into 1mL of complete medium or into a mixture of 800µL complete medium and 200µL collagenase stock, resulting in a final collagenase concentration of 2mg/mL. Slices were incubated at 37°C and 10% CO_2_ for 20 minutes. The medium was then removed, and slices were washed five times with complete medium to remove any remaining enzymatic activity. Following treatment, lymph node slices were either immediately fixed for subsequent antibody staining or imaged live. For live imagine, lymph node slices were transferred to glass-bottom dishes. A small dome of complete medium was added, a glass coverslip was gently placed over the tissue, and excess medium was removed from beneath the coverslip to secure the slice in place before imaging.

### Vibratome slices antibody staining

Lymph node slices were fixed in 1mL of Antigen Fix (DiaPath, P0016) on ice with gentle agitation for 1 hour. Samples were then washed once with ice-cold PBS and transferred to 500µL of 0.1M Tris-HCl buffer, pH 7.4 (prepared by dissolving 12.1g Tris base in distilled water, adjusting the pH with HCl, and bringing the volume to 1L). Sections were subsequently incubated in 500⍰ µL of IHC buffer (0.5% BSA, 2% Triton X-100, 0.1M Tris-HCl, pH 7.4, in distilled water; see Reagents List) on ice with gentle agitation for 20 minutes. Primary antibodies were vortexed and diluted in 300µL of IHC buffer, and slices were incubated in this solution on ice for 2 hours with gentle agitation. LN slices were washed twice for 15 minutes in 1mL of 0.1M Tris-HCl buffer on ice with agitation. Secondary antibodies were vortexed and together with DAPI were added to 300µL of IHC buffer, in which LN slices were incubated on ice for 2 hours with agitation. LN slices were washed twice for 15 minutes in 1mL of 0.1M Tris-HCl buffer on ice with agitation. The lymph node slices were mounted in Mowiol (see reagent list).

**Table 1.**
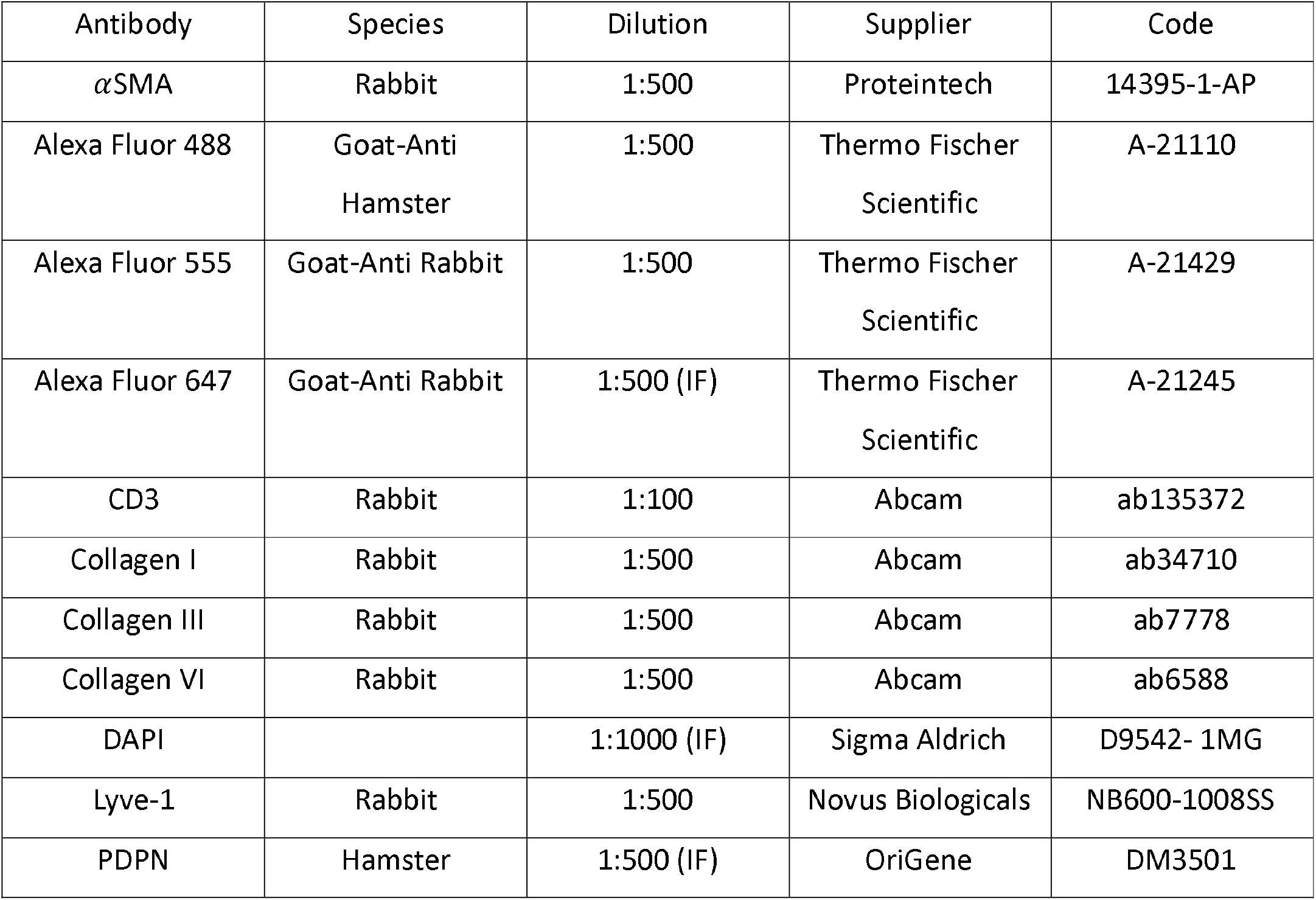
Antibodies used in the study.

| Antibody | Species | Dilution | Supplier | Code |
| --- | --- | --- | --- | --- |
| αSMA | Rabbit | 1:500 | Proteintech | 14395-1-AP |
| Alexa Fluor 488 | Goat-Anti<br>Hamster | 1:500 | Thermo Fischer<br>Scientific | A-21110 |
| Alexa Fluor 555 | Goat-Anti Rabbit | 1:500 | Thermo Fischer<br>Scientific | A-21429 |
| Alexa Fluor 647 | Goat-Anti Rabbit | 1:500 (IF) | Thermo Fischer<br>Scientific | A-21245 |
| CD3 | Rabbit | 1:100 | Abcam | ab135372 |
| Collagen I | Rabbit | 1:500 | Abcam | ab34710 |
| Collagen III | Rabbit | 1:500 | Abcam | ab7778 |
| Collagen VI | Rabbit | 1:500 | Abcam | ab6588 |
| DAPI |  | 1:1000 (IF) | Sigma Aldrich | D9542- 1MG |
| Lyve-1 | Rabbit | 1:500 | Novus Biologicals | NB600-1008SS |
| PDPN | Hamster | 1:500 (IF) | OriGene | DM3501 |

### Cryosectioning of lymph nodes

Lymph nodes used for cryosectioning, and immunofluorescence were fixed in 1mL of Antigen Fix (DiaPath, P0016) overnight at 4°C. The following day, lymph nodes were washed twice in 10mL of PBS and dehydrated overnight in 2mL of 30% sucrose (Sigma-Aldrich, S0389-500G) containing 0.01% (w/v) sodium azide (Sigma-Aldrich, 1066880100). The next day, lymph nodes were incubated for 1 hour in a 1:1 mixture of 30% sucrose and Tissue-Tek OCT compound (Sakura, 4583), then embedded in Tissue-Tek OCT in disposable base molds (Avantor, M475-2) and snap-frozen on dry ice. Frozen blocks were stored at −80°C until cryosectioning. Cryosectioning was performed using a Leica CM1950 cryostat. Sections were cut at a thickness of 15µm, mounted onto Superfrost™ microscope slides (Avantor, 631-0446) and stored at −80°C until further use.

### Live imaging of lymph node sections

Live lymph node sections were transferred to the bottom of a 35mm glass-bottomed dish (MatTek Corporation, P35G-1.5-20-C). A total of 50µL of RPMI 1640 medium supplemented with 1% ITS and 1% Penicillin-Streptomycin was added to each section, and a glass coverslip was gently placed on top. Excess medium was removed from beneath the coverslip to secure the tissue in place. Live imaging was performed using a Zeiss LSM 880 inverted multiphoton microscope. The imaging chamber was maintained at 37°C, and the stage-top incubator was set to deliver 10% CO_2_. A Plan Apochromat 40x oil objective was used for imaging.

### Confocal microscopy

Unless otherwise stated, all confocal microscopy was performed using a Zeiss LSM 900 inverted microscope. Images were acquired with 2-line averaging, a resolution of 1024⍰×⍰1024 pixels, and scan speed set from 6 to 8 depending on the sample. Fluorophore excitation and signal acquisition were carried out sequentially using bidirectional scanning. Fluorescence detection was performed using built-in GaAsP detectors. Regions of interest (ROIs) were selected manually, and z-stacks were configured accordingly. Z-step intervals ranged from 0.2µm to 0.7µm depending on sample type and thickness. Automatic tile stitching was performed using Zeiss ZEN software for tilescans.

### Laser Ablation Experiments

LN were sliced into 200µm sections using the vibratome as described previously and collected into complete media (Table 2.1). All laser ablations were performed on a Zeiss LSM 880 inverted multiphoton microscope equipped with a Chameleon Vision II Ti:Sa laser (Coherent), pulsed (<100 milliseconds), at 85-100% power and 760 nm wavelength. Laser ablations were carried out using a manually defined rectangle drawn across an FRC branch, away from the cell body, and a single z-plane was ablated. A single-channel time-lapse image sequence was acquired over 50 frames using a PMT detector, with a frame size of 512 × 512 pixels and a frame interval of 0.47491sec. One frame was acquired before laser ablation.

### Laser Ablation Analysis

Recoil of the FRC network following laser ablation was quantified using the MTrackJ plugin in Fiji. Displacement was measured as the change in distance between two points located on either side of the ablation site. Displacement-time curves were fitted to a Kelvin-Voigt viscoelastic model as previously described^27^. Initial recoil and the elasticity-viscosity ratio were extracted from the fitted parameters. Non-linear curve fitting was performed in GraphPad Prism 7 (GraphPad Software).

### Cellpose and Napari FRC network segmentation

FRC network segmentation was performed using Cellpose (v3.1.1.1) and Napari (v0.5.6). Binarised images of the FRC network were imported into Cellpose in .tif format and calibrated to obtain a suggested cell diameter. Cyto3 was then run to generate the initial segmentation, which was saved and subsequently imported into Napari together with the corresponding binarised image. The Cellpose output was converted to label format, and all manual corrections were performed in Napari.

### T-cell zone area and eccentricity quantification

T-cell zone areas were segmented as described in Section 2.1.12. Label masks were imported into Python, and the area and eccentricity of each segmented T-cell zone were extracted separately using the regionprops_table function from the scikit-image (skimage 0.25.0) library. Area measurements were converted from pixels to µm^2^ using the calibrated pixel size of the acquisition. For each region of interest (ROI), the mean area and mean eccentricity across all segmented T-cell zones were calculated, and these ROI-level means were compared across timepoints (Day 0, 5, 9, 14) using a Kruskal-Wallis test (see 2.1.14 Statistical analysis).

### ECM and FRC networks’ width analysis

All FRC and ECM thickness measurements were performed in Fiji (ImageJ) using the Local Thickness (masked, calibrated, silent) plugin on binary images of the network. This plugin calculates, at each point within the network, the diameter of the largest sphere that can be fitted entirely within the structure, and outputs a thickness histogram for each ROI.

### FRC-ECM colocalisation analysis

Colocalisation between the FRC network (PDPN) and ECM (Collagen I) was quantified in Fiji. The PDPN channel was thresholded to generate a binary mask, and the area of this initial PDPN network was measured. Collagen I channel was thresholded separately to generate a binary ECM mask, which was then dilated by 1 pixel (Process > Binary > Dilate, default settings: 1 iteration, count 1) uniformly across all ROIs to define a proximity zone around the collagen fibres. This dilated Collagen I mask was overlaid onto the PDPN mask, and the overlapping region was subtracted from the PDPN mask. The resulting image was re-thresholded, and the area of the remaining PDPN network was measured. The percentage of the FRC network lacking associated ECM was calculated as: % missing ECM = (remaining PDPN area / initial PDPN area) × 100.

### Integrated density measurement

For collagenase digestion experiments (Figure 4.2.1B), Collagen I, Collagen III and SHG signal intensities were quantified in Fiji using the Integrated Density measurement (Analyze > Measure). Regions of interest were drawn as large as possible within each field of view while excluding blood vessels, which showed disproportionately high signal intensity relative to the surrounding tissue. No background subtraction was performed prior to measurement, as signal was clean.

### PIV analysis

Particle image velocimetry (PIV) was performed in Fiji (Analyze > Optical Flow > PIV Analysis) to visualise relative tissue movement. As arrow colour and size scales were not standardised across analyses, PIV outputs are presented as representative images to illustrate qualitative patterns of movement only, and were not used for quantitative comparison between conditions.

### TWOMBLI analysis

Collagen I fibre curvature was quantified using the TWOMBLI (The Workflow Of Matrix BioLogy Informatics) FIJI macro, using default settings except for a curvature window of 10 pixels. Only the curvature output was used for downstream analysis and figure reporting; other TWOMBLI-derived metrics (e.g., fibre alignment, lacunarity, total fibre length) were not used.

### Patient data

Representative images of human axillary lymph node tissue from three patients were obtained from a previously described archival cohort of formalin-fixed paraffin-embedded (FFPE) samples collected through the Breast Cancer Immune, Drug and Gene (BRIDGE) Study (Research Ethics Committee Number: 24/NW/0079). Full details of cohort selection and clinicopathological characteristics have been reported previously (reference Jpath paper with whichever style requested by journal). For PDGFRβ immunohistochemistry, 3-μm FFPE sections were stained using a recombinant rabbit monoclonal anti-PDGFRβ antibody (clone RM303; Invitrogen MA5-33050) at 1:100 dilution on a Leica Bond III Autostainer.

### Statistical analysis

Where multiple regions of interest were sampled from the same lymph node, values were averaged to obtain a single measurement per lymph node prior to statistical analysis; lymph nodes, not individual regions of interest, were treated as the independent unit of analysis. For laser ablation experiments, each ablation was treated as an independent measurement, consistent with previous studies^3,18^; the number of ablations and independent experiments for each condition is reported in the corresponding figure legend.

Prior to hypothesis testing, each dataset was assessed for normality (Shapiro-Wilk test) and homogeneity of variance across groups (Levene’s test). Where these assumptions were met, differences between more than two groups were assessed by one-way or two-way ANOVA, as appropriate to the experimental design. Post-hoc correction was performed by Tukey’s multiple-comparisons test where all pairwise comparisons were of interest, by Dunnett’s multiple-comparisons test where multiple groups were compared against a single common control, or by Šídák’s multiple-comparisons test where only a pre-specified subset of pairwise comparisons was of interest (e.g., each timepoint relative to baseline); the specific post-hoc test used is indicated in the corresponding figure legend. Where normality and/or variance homogeneity were violated, the non-parametric Kruskal-Wallis test was used, followed by Dunn’s multiple-comparisons test with correction for the number of comparisons performed.

Two-group comparisons of independent samples were assessed by unpaired t-test (with Welch’s correction where variances were unequal) or Mann-Whitney U test, as appropriate; paired comparisons used a paired t-test or Wilcoxon matched-pairs signed-rank test. All two-group comparisons were two-sided.

For *in silico* data, each simulated condition comprised n = 3 independent simulations unless otherwise stated; comparisons between conditions were performed using the same statistical framework described above. Where the number of individual comparisons was too large to report in full in a figure legend, exact test statistics and adjusted p-values for all comparisons are provided in the corresponding Supplementary Table 1 or Table 2.

Data collection and analysis were not performed blinded to experimental conditions.

## Model Overview

All simulations were implemented using the Tyssue Python library (reference). We developed a two-dimensional vertex-based mechanical model of the fibroblastic reticular cell (FRC) network and associated extracellular matrix (ECM). T cell areas are represented as polygons whose vertices are connected by edges, and their vertex positions evolve under viscous, force-balanced dynamics, such that the tissue relaxes towards mechanical equilibrium. The model allows for continuous mechanical relaxation interspersed with discrete topological remodelling events, including edge collapse, vertex division, and ECM breakage. T1 transitions are not used.

### Computational model

We adapted a 2D vertex model using Tyssue Python library^18^. T cell areas are represented as polygons whose vertices are connected by edges, and their vertex positions evolve under viscous, force-balanced dynamics, such that the tissue relaxes towards mechanical equilibrium. T1 transitions are not used in the model setup.

The total energy of the system (*E*) is expressed as the sum of its effectors, or energy terms, including Cell Area Elasticity *E*_*AE*_, Length Elasticity *E*_*LE*_, and Line Tension *E*_*LT*_. More specifically, the total energy is a function of the vertex coordinates 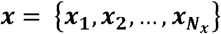 where *N*_*x*_ is the total vertex number.

#### Model 1

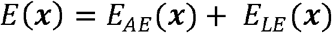

where,

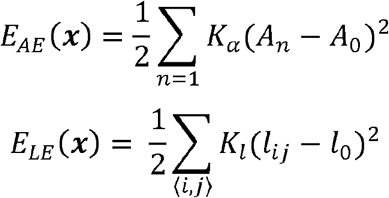

#### Model 2

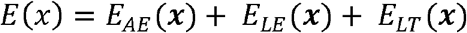

where,

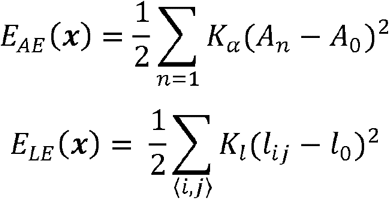

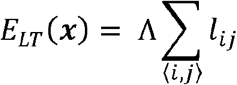

Here, *n= 1, …, N*_*c*_ is the T cell area index, and *l*_*ij*_ denotes the length of an edge connecting vertices *i* and *j*.

The term *E*_*AE*_ refers to the area elasticity energy, where *A*_*α*_ is the area of the T cell area, *n, A*_*0*_ is preferred area and *K*_*α*_ is area elasticity coefficient.

The term *E*_*LE*_ refers to the length elasticity energy, where *l*_*ij*_ is the length of the edge *(i, j), l*_*0*_ is preferred length and *K*_*l*_ is length elasticity coefficient.

The term *E*_*LT*_ refers to the line tension energy, where *l*_*ij*_ is the length of the edge *(i, j)*, and *Λ* is line tension coefficient.

Following the definition of the total mechanical energy *E ({x*_*i*_*})*, vertex motion is governed by overdamped force-balance dynamics. We assume that vertex velocities are proportional to the mechanical forces acting on them. Forces are defined as the negative gradient of the total energy with respect to vertex position,

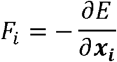

where *x*_*i*_ = (*x*_*i*_,*y*_*i*_) denotes the position of vertex *i*.

Under this assumption, the equation of motion for each vertex reads

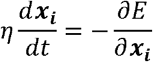

where *η> 0* is an effective friction coefficient that sets the timescale of mechanical relaxation as used in an explicit Euler method. Thus, the time dependent solved is written as follows:

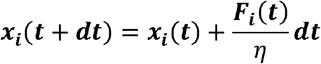

### Initial geometry

Simulations were initialised from a two-dimensional planar tissue generated using the Tyssue library. An approximately square patch of polygonal domains representing T cell areas was constructed using a Voronoi tessellation seeded on a regular lattice with controlled geometric noise (noise=0.2). The initial lattice spacing was uniform in both spatial directions, and positional noise was introduced to break perfect hexagonal symmetry. Boundary cells were trimmed by keeping only cells within the coordinate range [1, number of cells to keep] in both x and y direction.

### Model Parameter Sweep

A parameter sweep was conducted for both models. For Model 1, area elasticity (10–100), length elasticity (10–200), and viscosity (400–1500) were varied, with 20 linearly spaced values sampled for each parameter. For Model 2, area elasticity (1–100; 5 values), length elasticity (10–350; 8 values), viscosity (10–1500; 11 values), and line tension (1–1000; 18 values) were varied. Unlike Model 1, parameter values in Model 2 were sampled from predefined, non-uniform sets rather than linearly spaced intervals (values stated in **Parameter sweep**).

For each parameter combination, the model was initialised with the specified mechanical parameters and allowed to relax to an energy minimum. To assess how well each parameter combination reproduced *ex vivo* FRC network mechanics, *in silico* laser ablations were performed by selecting a random non-boundary edge and setting its length elasticity to zero in Model 1 and, in Model 2, both its length elasticity and line tension to zero, thereby mimicking laser ablation as performed in previous studies^27^. For each parameter combination, five laser ablations were performed consecutively on the same initialised model in Model 1, whereas three consecutive laser ablations were performed in Model 2, with energy minimisation and response recording carried out after each ablation.

The initial recoil and elasticity-viscosity ratio were then calculated using the same approach as for the *ex vivo* data. Initial recoil and elasticity-viscosity ratio (K) were obtained by fitting the post-ablation displacement curve to a nonlinear exponential recoil model, 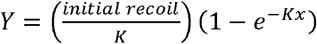. Mahalanobis distance analysis was used to quantify the similarity between simulated and *ex vivo* measurements, and the parameter combination with the smallest Mahalanobis distance in both models was selected for further analysis.

### Parameter sweep

#### Area Elasticity

- Model 1: 10, 20, 30, 40, 50, 60, 70, 80, 90, 100, 110, 120, 130, 140, 150, 160, 170, 180, 190, 200
- Model 2: 1, 10, 20, 50, 100

#### Length Elasticity

- Model 1: 10, 20, 30, 40, 50, 60, 70, 80, 90, 100, 110, 120, 130, 140, 150, 160, 170, 180, 190, 200
- Model 2: 10, 50, 100, 150, 200, 250, 300, 350

#### Viscosity

- Model 1: 400.0, 457.89, 515.79, 573.68, 631.58, 689.47, 747.37, 805.26, 863.16, 921.05, 978.95, 1036.84, 1094.74, 1152.63, 1210.53, 1268.42, 1326.32, 1384.21, 1442.11, 1500.0
- Model 2: 10.0, 100.0, 175.6, 341.1, 506.7, 672. 2, 837. 8, 1003.3, 1168. 9, 1334.4, 1500.0

#### Line Tension

- Model 2: 1, 2, 4, 6, 10, 16, 24, 39, 61, 96, 152, 241, 380, 600, 700, 800, 900, 1000

### Collagenase Simulation

For Model 1, the FRC network was initialised with parameters selected from the parameter sweep. The Length Elasticity term was partitioned between *K*_*FRC*_ and *K*_*ECM*_ across a range of FRC:ECM ratios: 1:1, 2:1, 3:1, 4:1, 5:1, 1:2, 1:3, 1:4, 1:5, 1:6, 1:7, 1:8, 1:9, and 1:10. For each ratio, following relaxation to a minimum-energy state, *K*_*ECM*_ *=* 0 to simulate collagenase treatment, and the network was subsequently allowed to relax to a new minimum-energy state. Upon reaching this new equilibrium, 5 laser ablations were performed on each relaxed network by severing a randomly selected interior edge (setting its Length Elasticity to zero), and recoil dynamics were recorded over 50 time units until complete energy minimisation. The initialisation process and laser ablations were repeated 3 times independently for each FRC:ECM ratio tested (15 total ablations per ratio).

For Model 2, the FRC network was initialised with parameters selected from the parameter sweep. The tissue was then relaxed to a minimum-energy state. To simulate complete collagenase-mediated ECM degradation, we set *K*_*ECM*_ = 0, effectively setting the Length Elasticity of all edges to zero, and the tissue was allowed to relax to a new equilibrium. Subsequently, 5 laser ablations were performed per independently initialised tissue patch by severing a randomly selected interior edge (setting *K*_*ECM*_ *= 0* and *Λ*_*FRC*_ *= 0*), with recoil dynamics recorded over 100 time units until complete energy minimisation. The initialisation and ablation process were repeated 3 times (15 total ablations).

### Adding a vertex in the middle of a chosen edge to model FRC division

To subdivide an existing edge, we insert a new vertex at the midpoint of the chosen edge. The procedure is as follows:

**Step 1:** The source and target vertices of the chosen edge are retrieved from the edge DataFrame. *Step 2:* Calculate midpoint coordinates. The coordinates of the new vertex are computed as the average of the source and target vertex coordinates: (x_mid, y_mid) = ((x_1_ + x_2_)/2, (y_1_ + y_2_)/2).

**Step 3:** A new vertex is added to the vertex DataFrame at the calculated midpoint. All mechanical properties (viscosity, activity status, etc.) are copied from the source vertex.

**Step 4:** Both the original directed edge (source → target) and its opposite half-edge (target → source) are removed from the edge DataFrame.

**Step 5:** Four directed edges are created to maintain proper topology: source → new_vert, new_vert → source, new_vert → target, and target → new_vert. The mechanical properties of these new edges are copied from the original edge.

**Step 6:** The two edges travelling in the original direction (source → new_vert and new_vert → target) inherit the face of the original edge. The two edges travelling in the opposite direction inherit the face of the opposite edge.

**Step 7:** Geometry is recalculated, topology is reset, and the system is allowed to relax by solving the equations of motion for 40 time units using Euler integration.

### Division by adding a vertex in a random place on a chosen edge

Division by adding a vertex at a random location along a chosen edge follows the same logic as adding a vertex at the midpoint, except the coordinates of the new vertex are calculated differently. Therefore, all the steps from **Adding a vertex in the middle of a chosen edge to model FRC division** are the same except for Step 2, where coordinates of the new vertex are calculated as follows:

**Step 2 (modified)**: Instead of placing the new vertex at the midpoint, we first sample a random fraction, f, uniformly from [0, 1]. The new vertex coordinates are then computed as a linear interpolation between the source and target: (x_new, y_new) = (x_1_ + f·(x_2_ - x_1_), y_1_ + f·(y_2_ - y_1_)). When f = 0.5, this reduces to the midpoint case. All remaining steps (vertex creation, edge deletion, creation of four new edges, face assignment, geometry update, and relaxation) are identical to the midpoint division procedure

### FRC Division by splitting a vertex

To model cell division by vertex splitting, we implemented an operation that divides a single vertex into two distinct vertices connected by a newly created edge. The procedure is as follows:

**Step 1**: Create a new vertex. A new vertex is created at a random location within a specified distance from the original vertex. The random direction is sampled uniformly from 0 to 2π. All mechanical properties of the new vertex are copied from the original vertex.

**Step 2**: Create a pair of opposite edges between the original vertex and the new vertex. The mechanical properties of these edges are copied from an edge connected to the original vertex, ensuring that the new edges inherit the same material properties as existing edges. The face assignment to these edges is NaN at this stage.

**Step 3**: Reassign surrounding edges. All edges connected to the original vertex are evaluated to determine whether they should remain attached to the original vertex or be reassigned to the new vertex. The assignment is based on Euclidean distance: each edge is reassigned to whichever vertex (original or new) is closer to the edge’s other endpoint.

**Step 4**: Identify open faces. After reassignment, some faces become “open,” meaning their edge chains no longer form closed loops, since the new pair of opposite edges has not been assigned a face. We identify open faces by traversing each face’s edge sequence and checking whether the first and last vertices match. Only open faces that touch either the original or new vertex are considered for further processing; exactly two such faces are expected (one on each side of the newly created edge).

**Step 5**: Assign faces to new edges. For each open face, we determine the missing edge required to close the loop by identifying the start and end vertices (where the count of incoming and outgoing edges is unbalanced). The newly created edge that matches the required direction is assigned to that face.

**Step 6**: Verify topological consistency. We verify that both faces now form closed directed cycles by checking that (i) the face forms a single, unbroken ring where you can start at any vertex and follow the arrows all the way around without getting stuck or hitting a dead end, and (ii) a complete walk following edge directions returns to the starting vertex after traversing all edges. If verification fails, the face assignments are swapped and rechecked; if still invalid, the attempt is aborted.

Failure recovery. If any step fails (e.g., incorrect number of open faces, failed closure verification), the tissue state is rolled back to the backup copies, and a new attempt is made with a different placement of the new vertex. After three failed attempts, the operation returns without modifying the tissue.

### FRC apoptosis through single edge contraction

The single edge collapse operation merges two adjacent vertices connected by a specified edge into a single vertex. First, the edge to be collapsed is identified (see **Model simulations of homeostasis with FRC division and apoptosis**) and the coordinates of its two endpoint vertices are noted. A midpoint location between these two vertices is calculated. A new vertex is inserted at this midpoint. The two original vertex IDs are then replaced with the new vertex ID in all edges (both as source and target). The original edge and its parallel edge are removed, followed by deletion of the two original vertices. Any self-loop edges (source equals target) and duplicate edges (identical source-target pairs) are removed. The geometry is updated, energy is recomputed, and the system is allowed to relax into an energy minimum state.

### FRC apoptosis by two edge contraction

An initial edge is selected for contraction based on a selection criterion (see **Model simulations of homeostasis with FRC division and apoptosis**). Single-edge contraction is then performed as described above, resulting in the generation of a new vertex at the collapse site and all connected neighbouring edges to it are identified. A neighbouring edge is subsequently selected at random and collapsed using the same single-edge contraction procedure, thereby simulating the sequential contraction of two adjacent edges. Finally, geometry is updated, energy is recomputed, and the system is allowed to relax into an energy minimum state.

### Contraction of all edges surrounding a vertex

A vertex is selected for contraction (see **Model simulations of homeostasis with FRC division and apoptosis**) and all edges connected to it are identified. The vertices located at the opposite ends of these connected edges (the surrounding vertices) are identified. All edges connected to the original vertex are deleted. Every edge connected to a surrounding vertex, is reconnected to the original vertex. The surrounding vertices are then deleted. Any invalid edges resulting from the operation (such as self-loops or duplicate edges) are removed. Finally, geometry is updated, energy is recomputed, and the system is allowed to relax into an energy minimum state.

### Model simulations of homeostasis with FRC division and apoptosis

Model was initialised with 40 T cell areas arranged in a hexagonal lattice as described in Model Setup. Boundary vertices were immobilised with high viscsocity, mimicking lymph node capsule. Homeostasis simulations alternated between edge contraction (apoptosis) and vertex division (cell division) for 500 cycles, resulting in 500 apoptotic events and 500 division events. For each cycle, an interior edge (at least one layer from the boundary) was selected using one of three criteria: random selection, shortest edge length, or longest edge length. The selected edge was contracted to its midpoint, followed by 40 time units of mechanical relaxation. Subsequently, an interior vertex was selected using one of three criteria: random selection, smallest total attached edge length, or largest total attached edge length. The selected vertex was divided by creating a new vertex at a random nearby location as described in Vertex Division (Split Vertex), followed by another 40 time units of relaxation. After 500 cycles, the resulting network’s geometry and topology were compared to *ex vivo* FRC network data.

### Network Expansion Simulations

A model of 40 × 40 T-cell areas was initialised using the best-fit parameters identified from the parameter sweep, outer vertices were immobilised by assigning a high viscosity (2000000), mimicking the mechanical constraint imposed by the lymph node capsule under homeostatic conditions. The model was first relaxed for 300 timesteps (Δt = 1) to allow the tissue to reach an energy minimum. Prior to expansion, the boundary was relaxed to the same extent in all simulations by setting the viscosity of outer vertices to 10,000 and reducing the line tension of boundary edges to 300. Following equilibration, parameters of interest were perturbed in isolation or in combination to probe their effects on tissue expansion (Supplementary Table 3). The perturbed tissue was then simulated for a further 15,000 timesteps, with energy minimisation performed every 500 timesteps. All expansion simulations were performed in triplicate using independently generated and relaxed cell maps.

### Simulating spatial patterns of mechanical pathology

All simulations of localised mechanical pathology were initiated from the same three FRC network checkpoints at simulation step 3,500 during ‘normal’ lymph node expansion, ensuring that each perturbation condition began from an identical network geometry and mechanical state. The checkpoint was generated using the expansion model described above, incorporating increased T-cell preferred area, reduced FRC line tension and mechanically regulated FRC division.

To model spatially heterogeneous mechanical pathology, patches were seeded at random positions within the FRC network and expanded until the specified proportion of network branches had been incorporated. Perturbations were applied to 10%, 30% or 50% of network branches and distributed across 1, 10 or 50 patches. As patches expanded, neighbouring patches were permitted to merge, such that the final spatial organisation emerged from the randomly selected seed positions. Both half-edges corresponding to each selected branch were assigned the same perturbation to preserve branch-level mechanical consistency. FRC hypercontractility was simulated by multiplying the line tension of selected branches by 1.2 or 1.5, whereas ECM stiffening was simulated by multiplying the ECM length elasticity of selected branches by 3 or 5. Following assignment of the mechanical perturbations, network topology was reset before simulations were continued.

Each perturbed network was subsequently simulated for an additional 12,000 time steps, from step 3,500 to step 15,500, using 500-step remodelling cycles. At each cycle, 0.05% of internal edges were randomly selected for collapse to represent FRC apoptosis, followed by 10 relaxation steps. FRC division was then implemented through vertex splitting (see **FRC Division by splitting a vertex**) for internal vertices whose summed connected edge lengths exceeded a threshold of 3.5au. Following division, the network underwent 500 mechanical relaxation steps before the next remodelling cycle.

Network states and corresponding visualisations were saved every 500 simulation steps for subsequent analysis.

## Code availability

Code is available at https://github.com/vlachina/LymphNode.

## References

1. Mehrdad Matloubian et al. Lymphocyte egress from thymus and peripheral lymphoid organs is dependent on S1P receptor 1. Nature (2004).

2. Assen, F. P. et al. Multitier mechanics control stromal adaptations in the swelling lymph node. Nat. Immunol. 23, 1246–1255 (2022).

3. Horsnell, H. L. et al. Lymph node homeostasis and adaptation to immune challenge resolved by fibroblast network mechanics. Nat. Immunol. 23, 1169–1182 (2022).

4. Najibi, A. J. et al. Durable lymph-node expansion is associated with the efficacy of therapeutic vaccination. Nat. Biomed. Eng. https://doi.org/10.1038/s41551-024-01209-3 (2024) doi:10.1038/s41551-024-01209-3.

5. Wyatt, T., Baum, B. & Charras, G. A question of time: Tissue adaptation to mechanical forces. Current Opinion in Cell Biology vol. 38 68–73 Preprint at 10.1016/j.ceb.2016.02.012 (2016).

6. Mandl, J. N. et al. Quantification of lymph node transit times reveals differences in antigen surveillance strategies of naïve CD4+ and CD8+ T cells. Proc. Natl. Acad. Sci. U. S. A. 109, 18036–18041 (2012).

7. Acton, S. E. et al. Dendritic cells control fibroblastic reticular network tension and lymph node expansion. Nature 514, 498–502 (2014).

8. Kaldjian, E. P., Gretz, J. E., Anderson, A. O., Shi, Y. & Shaw, S. Spatial and Molecular Organization of Lymph Node T Cell Cortex: A Labyrinthine Cavity Bounded by an Epithelium-like Monolayer of Fibroblastic Reticular Cells Anchored to Basement Membrane-like Extracellular Matrix. International Immunology vol. 13 (2001).

9. Sixt, M. et al. The conduit system transports soluble antigens from the afferent lymph to resident dendritic cells in the T cell area of the lymph node. Immunity 22, 19–29 (2005).

10. Humphrey, J. D., Dufresne, E. R. & Schwartz, M. A. Mechanotransduction and extracellular matrix homeostasis. Nature Reviews Molecular Cell Biology vol. 15 802–812 Preprint at 10.1038/nrm3896 (2014).

11. Genovese, L. & Brendolan, A. Lymphoid tissue mesenchymal stromal cells in development and tissue remodeling. Stem Cells International vol. 2016 Preprint at 10.1155/2016/8419104 (2016).

12. Du Bois, H., Heim, T. A. & Lund, A. W. Tumor-Draining Lymph Nodes: At the Crossroads of Metastasis and Immunity. Sci. Immunol vol. 6 https://www.science.org (2021).

13. Hayakawa, M., Kobayashi, M. & Hoshino, T. Direct Contact between Reticular Fibers and Migratory Cells in the Paracortex of Mouse Lymph Nodes: A Morphological and Quantitative Study. Arch. Histol. Cytol vol. 51 (1988).

14. Sobocinski, G. P. et al. Ultrastructural localization of extracellular matrix proteins of the lymph node cortex: Evidence supporting the reticular network as a pathway for lymphocyte migration. BMC Immunol. 11, (2010).

15. Gregory, J. L. et al. Infection Programs Sustained Lymphoid Stromal Cell Responses and Shapes Lymph Node Remodeling upon Secondary Challenge. Cell Rep. 18, 406–418 (2017).

16. Kim, J. Il et al. CRISPR/Cas9-mediated knockout of Rag-2 causes systemic lymphopenia with hypoplastic lymphoid organs in FVB mice. Lab. Anim. Res. 34, 166–175 (2018).

17. Martinez, V. G. et al. Fibroblastic Reticular Cells Control Conduit Matrix Deposition during Lymph Node Expansion. Cell Rep. 29, 2810–2822.e5 (2019).

18. Theis, S., Suzanne, M. & Gay, G. Tyssue: an epithelium simulation library. J. Open Source Softw. 6, 2973 (2021).

19. Acton, S. E. et al. Podoplanin-Rich Stromal Networks Induce Dendritic Cell Motility via Activation of the C-type Lectin Receptor CLEC-2. Immunity 37, 276–289 (2012).

20. Astarita, J. L. et al. The CLEC-2-podoplanin axis controls the contractility of fibroblastic reticular cells and lymph node microarchitecture. Nat. Immunol. 16, 75–84 (2015).

21. Ferris, R. L., Lotze, M. T., Leong, S. P. L., Hoon, D. S. B. & Morton, D. L. Lymphatics, lymph nodes and the immune system: Barriers and gateways for cancer spread. in Clinical and Experimental Metastasis vol. 29 729–736 (2012).

22. Llewellyn, A. M. et al. Topological analysis of the human lymph node reticular network predicts outcome in breast cancer. Journal of Pathology https://doi.org/10.1002/path.70065 (2026) doi:10.1002/path.70065.

23. Hinz, B., Celetta, G., Tomasek, J. J., Gabbiani, G. & Chaponnier, C. Alpha-Smooth Muscle Actin Expression Upregulates Fibroblast Contractile Activity. Molecular Biology of the Cell vol. 12 (2001).

24. Klingberg, F., Hinz, B. & White, E. S. The myofibroblast matrix: Implications for tissue repair andfibrosis. Journal of Pathology vol. 229 298–309 Preprint at 10.1002/path.4104 (2013).

25. Miwa, H. & Era, T. Generation and characterization of PDGFRα-GFPCreERT2 knock-In mouse line. Genesis 53, 329–336 (2015).

26. Quétier, I. et al. Knockout of the PKN Family of Rho Effector Kinases Reveals a Non-redundant Role for PKN2 in Developmental Mesoderm Expansion. Cell Rep. 14, 440–448 (2016).

27. Liang, X., Michael, M. & Gomez, G. Measurement of Mechanical Tension at cell-cell junctions using two-photon laser ablation. Bio. Protoc. 6, (2016).

