## Extended data figures for "The Extracellular Matrix Regulates Tissue Mechanics to Enable Cyclic Lymph Node Remodelling for Sustained Immunity"

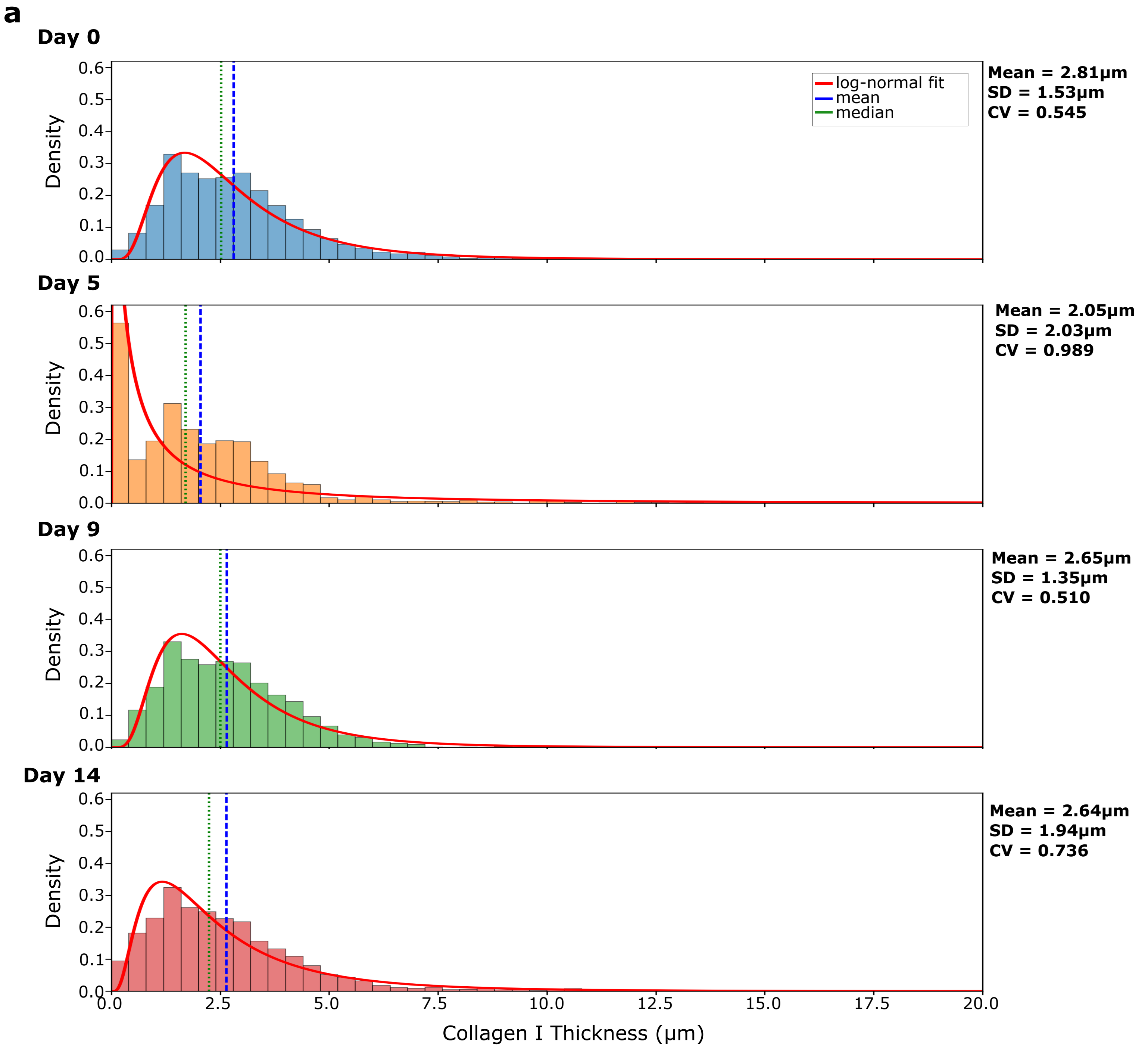


**Extended Data Figure 1 | Collagen I thickness distribution across immunisation. a**, Collagen I thickness distributions by day (Day 0, 5, 9, 14), with log-normal fits (red line). Blue dashed line = mean; green dotted line = median. Standard deviation (SD) and coefficient of variation (CV) shown next to plots. Data were collected from n = 26-31 ROIs per day, from N = 5-7 independent lymph nodes. Data in support of Figure 1.

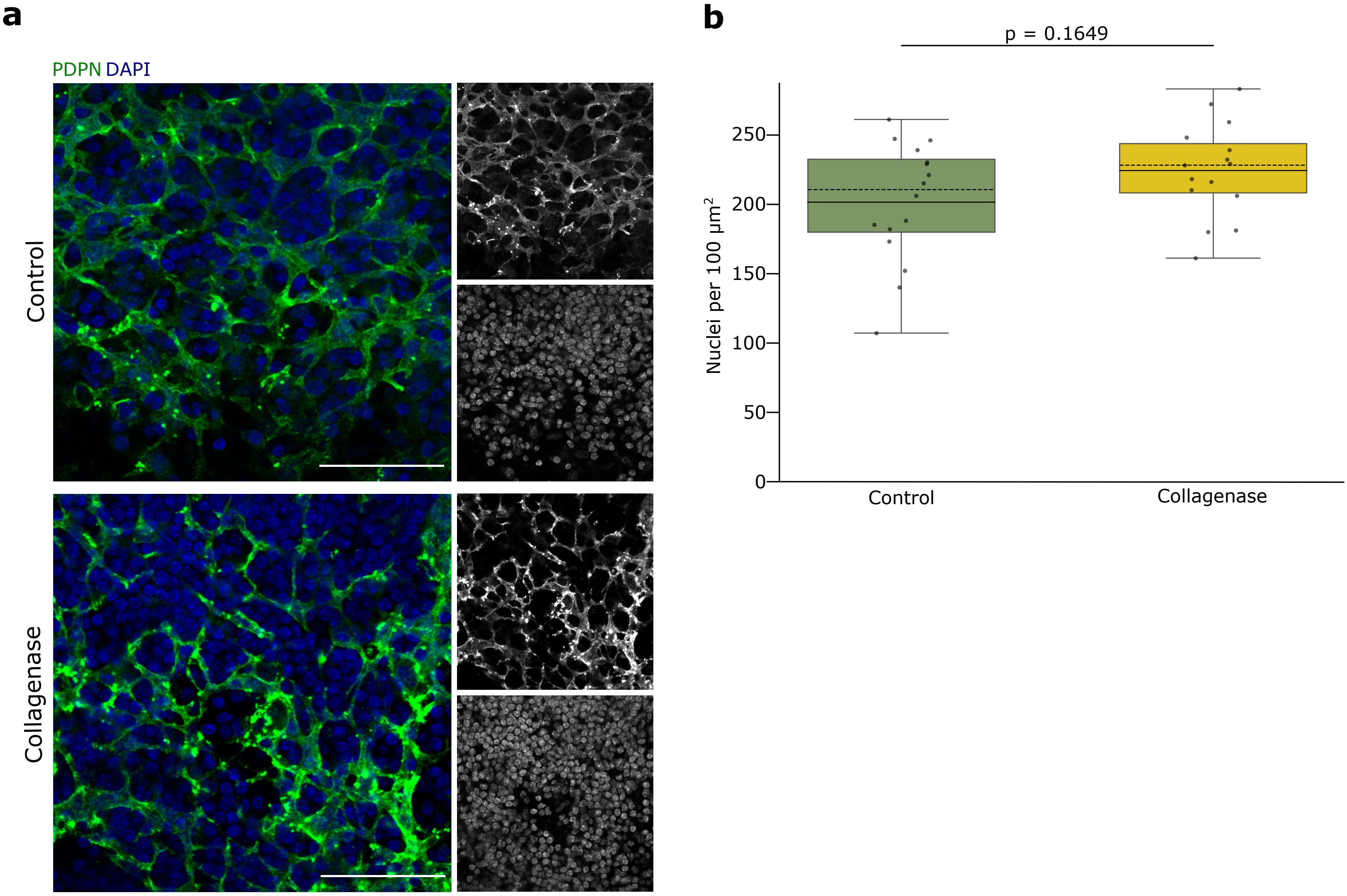


**Extended Data Figure 2 | Nuclei density in control and collagenase-treated lymph nodes.**
**a**, Representative immunofluorescence staining of FRC network (PDPN, green) and cell nuclei (DAPI, blue). Scale bar, 50μm. **b**, Quantification of nuclei density per 100μm². Box plots show distribution with box = IQR; whiskers = 1.5×IQR; mean = solid line; median = dashed line. Each dot is an individual ROI (n = 2-5 per LN); statistical analysis performed on LN means (N = 4 paired LNs). Statistics: paired t-test, p value shown. Data in support of Figure 2.

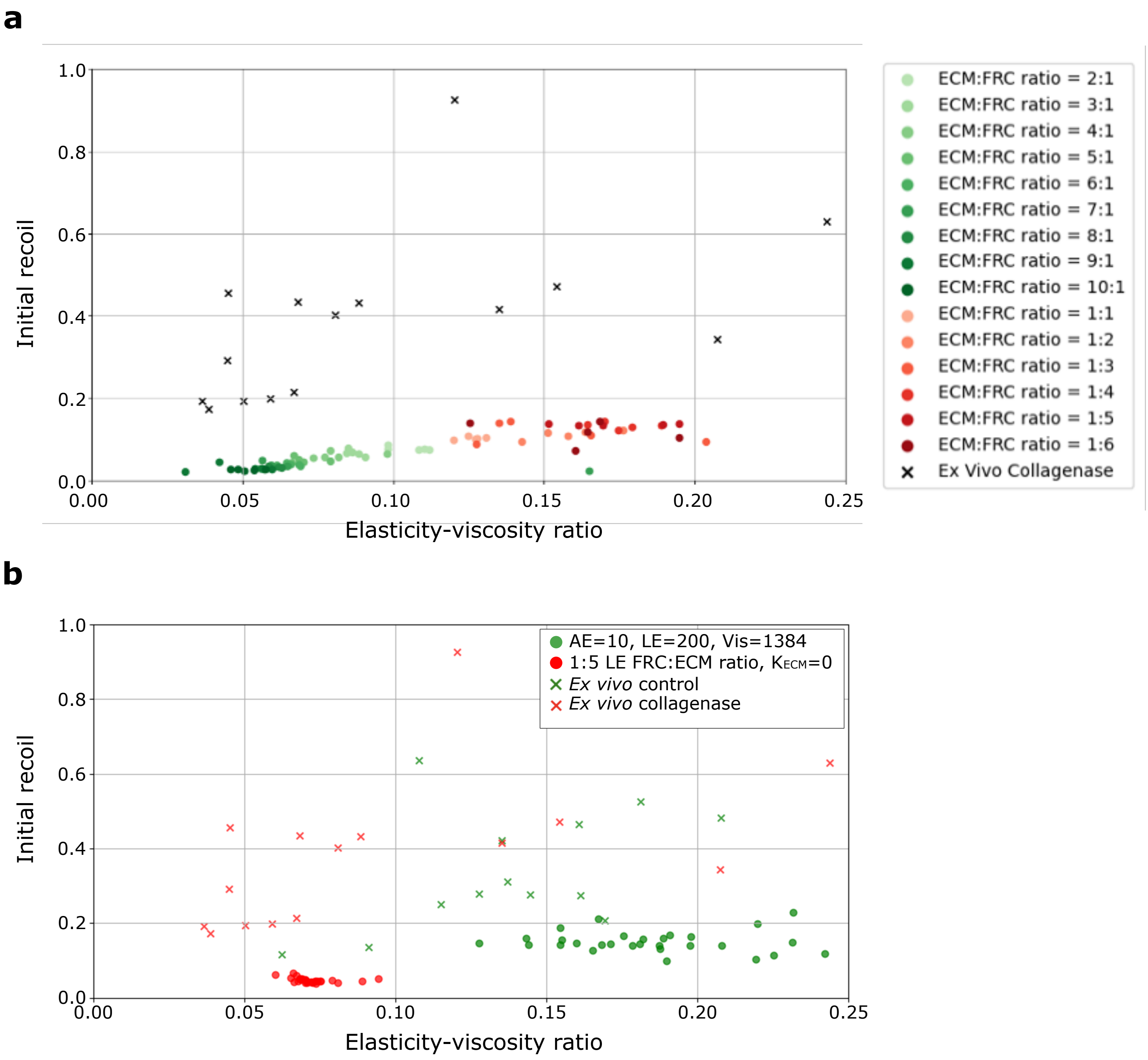


**Extended Data Figure 3 | Selection of the ECM:FRC length elasticity ratio in Model 1**. **a**, Length elasticity was partitioned between extracellular matrix (ECM) and cellular (FRC) components across ECM:FRC ratios ranging from 1:6 (red) to 10:1 (green) in Model 1. ECM length elasticity was subsequently set to zero to simulate collagenase treatment, followed by *in silico* laser ablations (N = 3 per ratio, from 3 independently initialised networks). The resulting initial recoil and elasticity-viscosity ratio were compared with *ex vivo* collagenase laser ablation measurements (black crosses; n = 16, from N = 4 independent experiments). An ECM:FRC ratio of 5:1 (equivalently, FRC:ECM = 1:5) most closely recapitulated the *ex vivo* collagenase response. **b**, To validate this parameterisation, an expanded set of 25 *in silico* laser ablations (n = 5 per network, from N = 5 independently initialised networks) was performed using the selected ratio under both control (green circles) and simulated collagenase conditions (red circles). These simulations reproduced the separation between control and collagenase observed in the *ex vivo* laser ablation experiments (control, green crosses, n = 13; collagenase-treated, red crosses, n = 16; both from N = 4 independent experiments). Data in support of Figure 4.

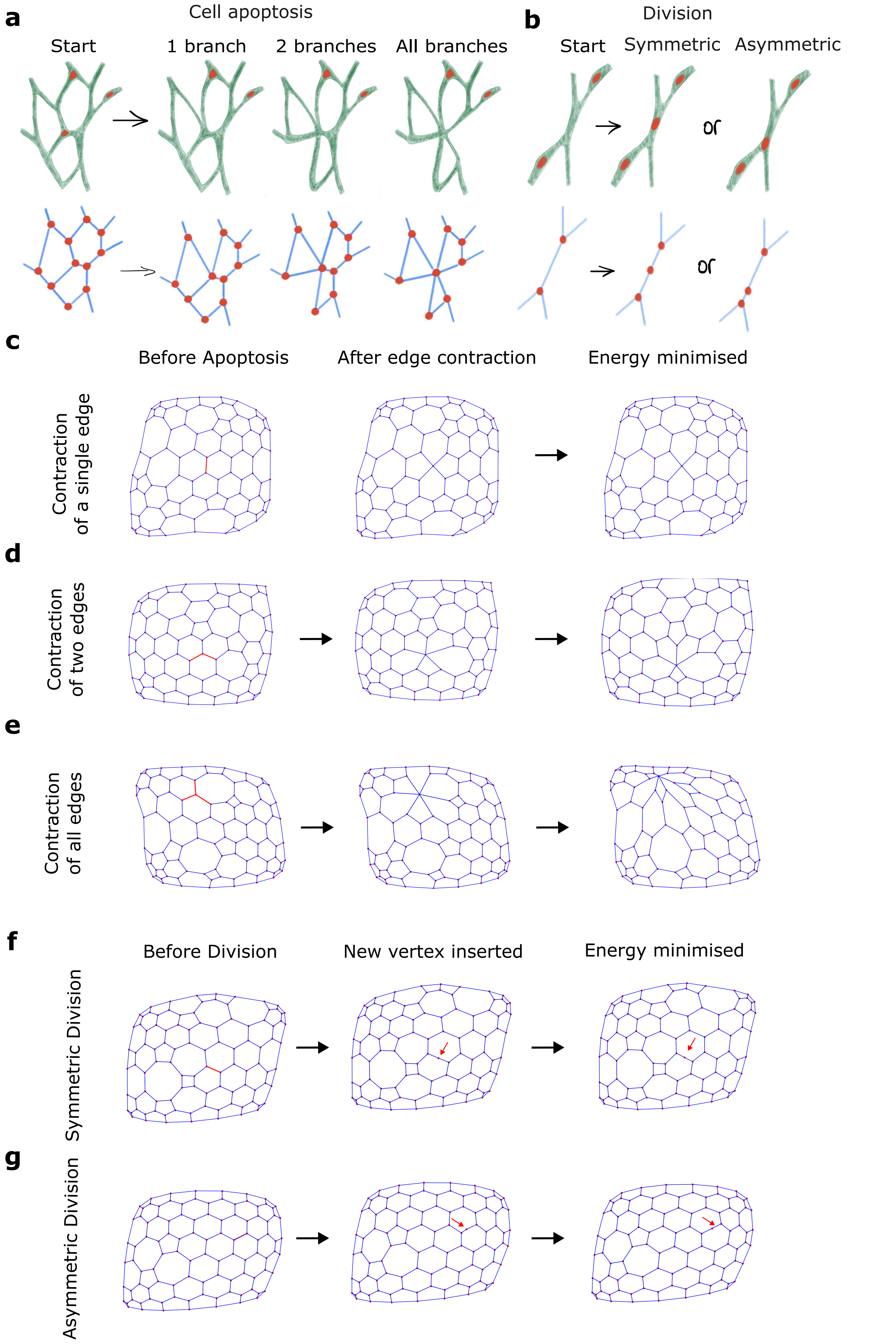


**Extended Data Figure 4 | Investigating different mechanisms of FRC division and apoptosis in the model. a,b**, Schematic of cell apoptosis and cell division mechanisms tested in the model. **c-e,** Comparison of three implementations of apoptosis in the vertex model: contraction of a single edge (**c**), contraction of two edges (**d**), or contraction of all edges surrounding a chosen vertex (**e**). **f,g**, Comparison of cell division implemented by inserting a new vertex in the middle of a chosen edge (**f**) or at a random point in the chosen edge (**g**). Data in support of Figure 5.

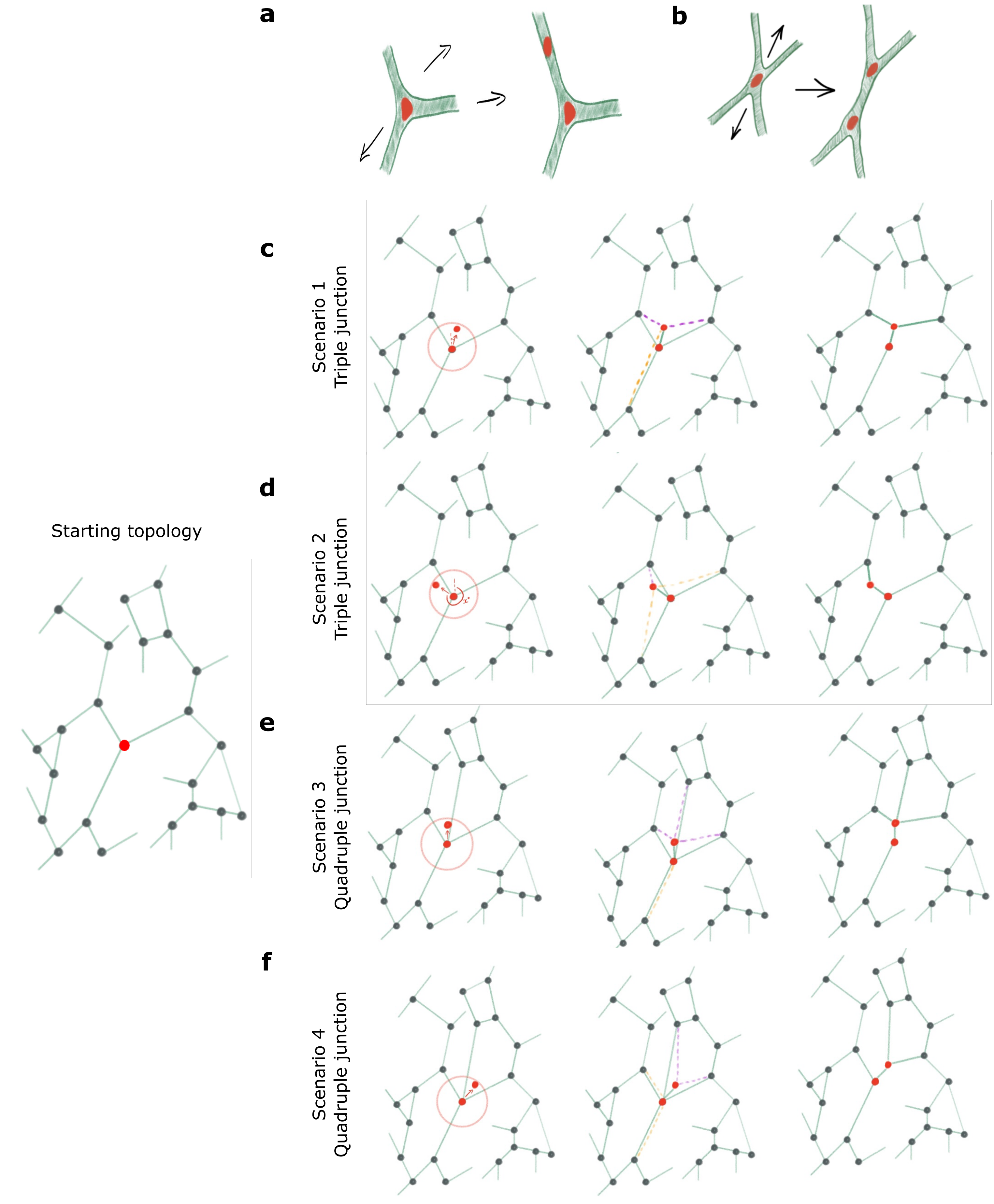


**Extended Data Figure 5 | Mechanism of division by vertex splitting in the model. a,b**, A schematic of cell dividing by vertex splitting in a triple and quadruple junction, respectively. **c-f**, Diagrams of the vertex splitting mechanism for different junction types, shown for a shared starting network topology (far left). A vertex is selected for division (red circle connected to the network), and a new vertex is inserted within a defined radius (red circle outside the network). Edges connected to the original vertex are reassigned based on proximity: edges closer to the new vertex are transferred to the new vertex (purple), while edges closer to the original vertex are retained (yellow). Example outcomes for division of tri- (**c,d**) and quadri- (**e,f**) junctions shown. Data in support of Figure 5.

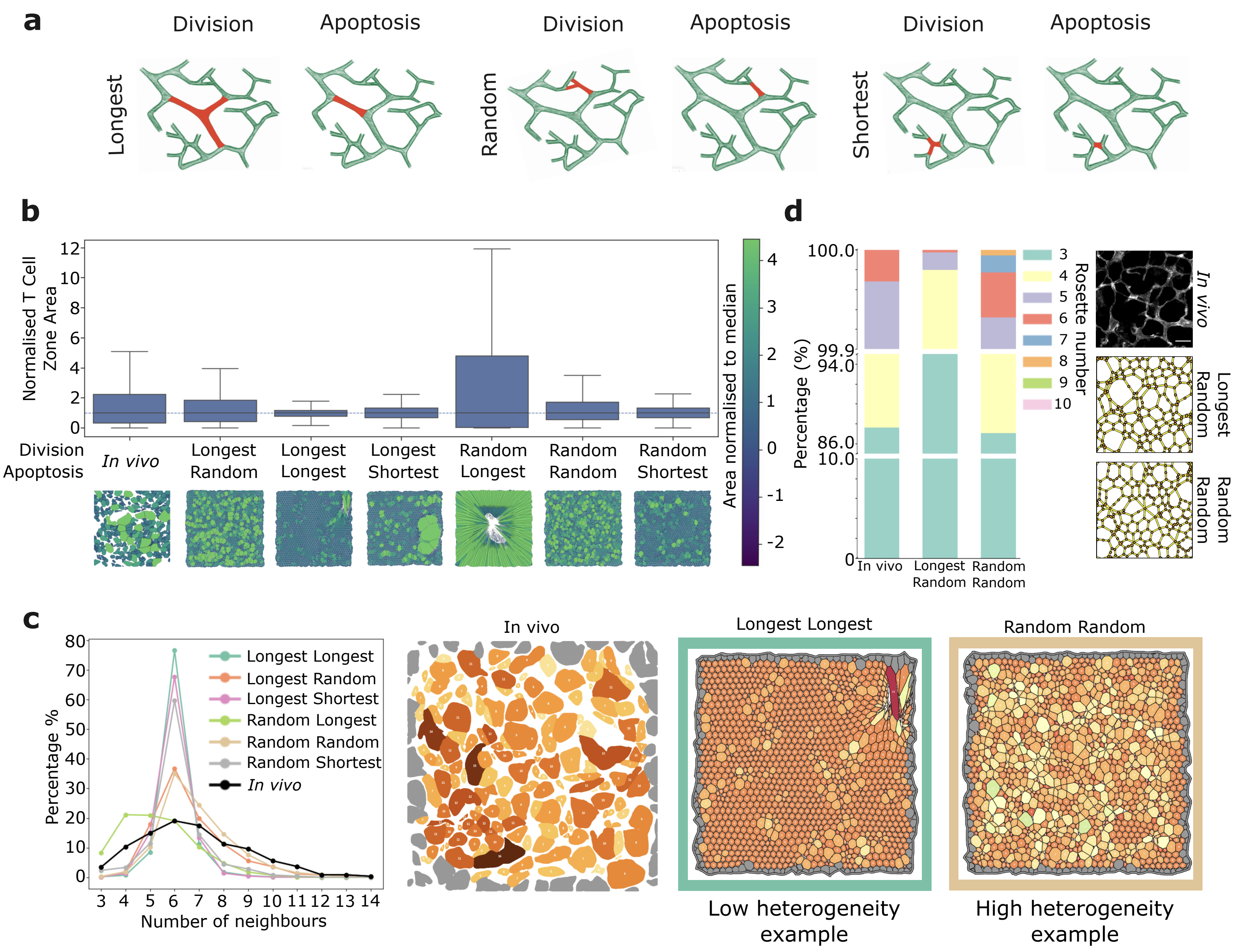


**Extended Data Figure 6 | Choosing rules for FRC division and apoptosis in the model.**
**a**, Schematic of the rules tested for selecting which FRC branch undergoes division or apoptosis: the longest, shortest, or a randomly selected branch. Division by the shortest branch was excluded from panels **b-d**, as it consistently divided the same shortest branch in a loop rather than distributing division across the network. **b**, Normalised T cell zone area distributions generated by each division-apoptosis strategy (N = 3 simulations per strategy, from the same 3 independently initialised networks across strategies), compared with *in vivo* measurements at day 0 (N = 2 LNs, n = 9 ROIs). Representative tissue snapshots are shown below each condition. Colour indicates area normalised to median. **c**, Distribution of cell neighbour number for each division-apoptosis strategy (as in **b**) compared with the *in vivo* tissue at day 0 (N = 2 LNs, n = 8 ROIs). Representative tissue images, coloured by T cell zone neighbour number, are shown for the *in vivo* tissue and two simulation conditions illustrating low (Longest Longest) and high (Random Random) heterogeneity. **d**, Rosette number distribution for the *in vivo* tissue at day 0 (N = 2 LNs, n = 8 ROIs) and the two best-performing simulation conditions from **b** and **c** (Longest Random and Random Random). Representative examples of rosette organisation are shown alongside each condition (*in vivo* scale bar, 10µm). Data in support of Figure 5.

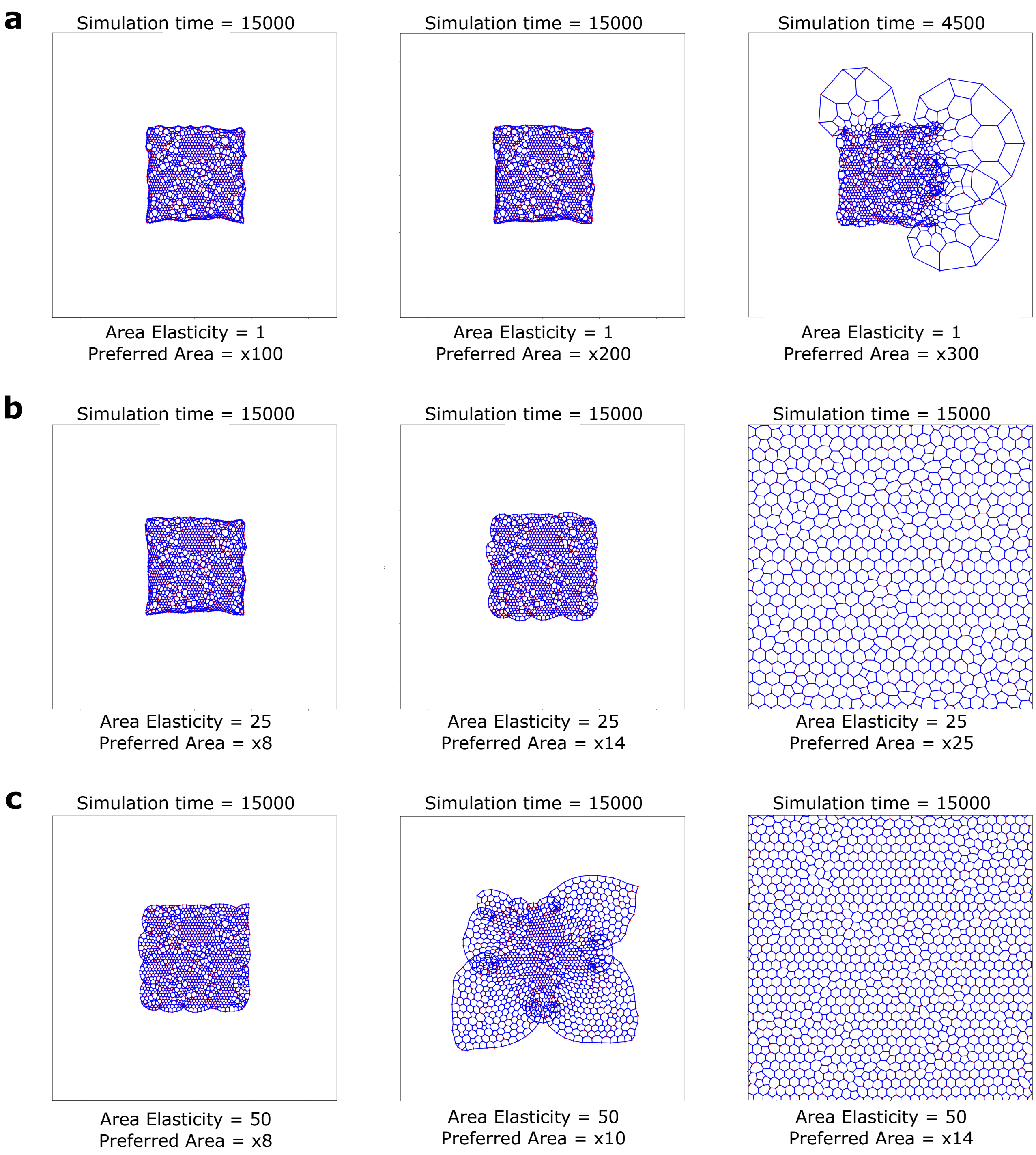


**Extended Data Figure 7 | Investigating whether pressure alone can drive lymph node expansion.** **a**, Representative simulations with Area Elasticity = 1 and increasing preferred area multiplier, illustrating the transition from limited expansion to numerical instability at large preferred areas; the simulation at x300 preferred area terminated early (t = 4500 vs. 15000 for all other conditions) due to this instability. **b**, Representative simulations with Area Elasticity = 25 across increasing preferred area multipliers, showing the transition from limited expansion to uniform tissue growth. **c**, Representative simulations with Area Elasticity = 50 across increasing preferred area multipliers, illustrating the effect of increasing preferred area on tissue morphology and expansion. Each condition reflects N = 3 independently initialised simulations. Data in support of Figure 5.

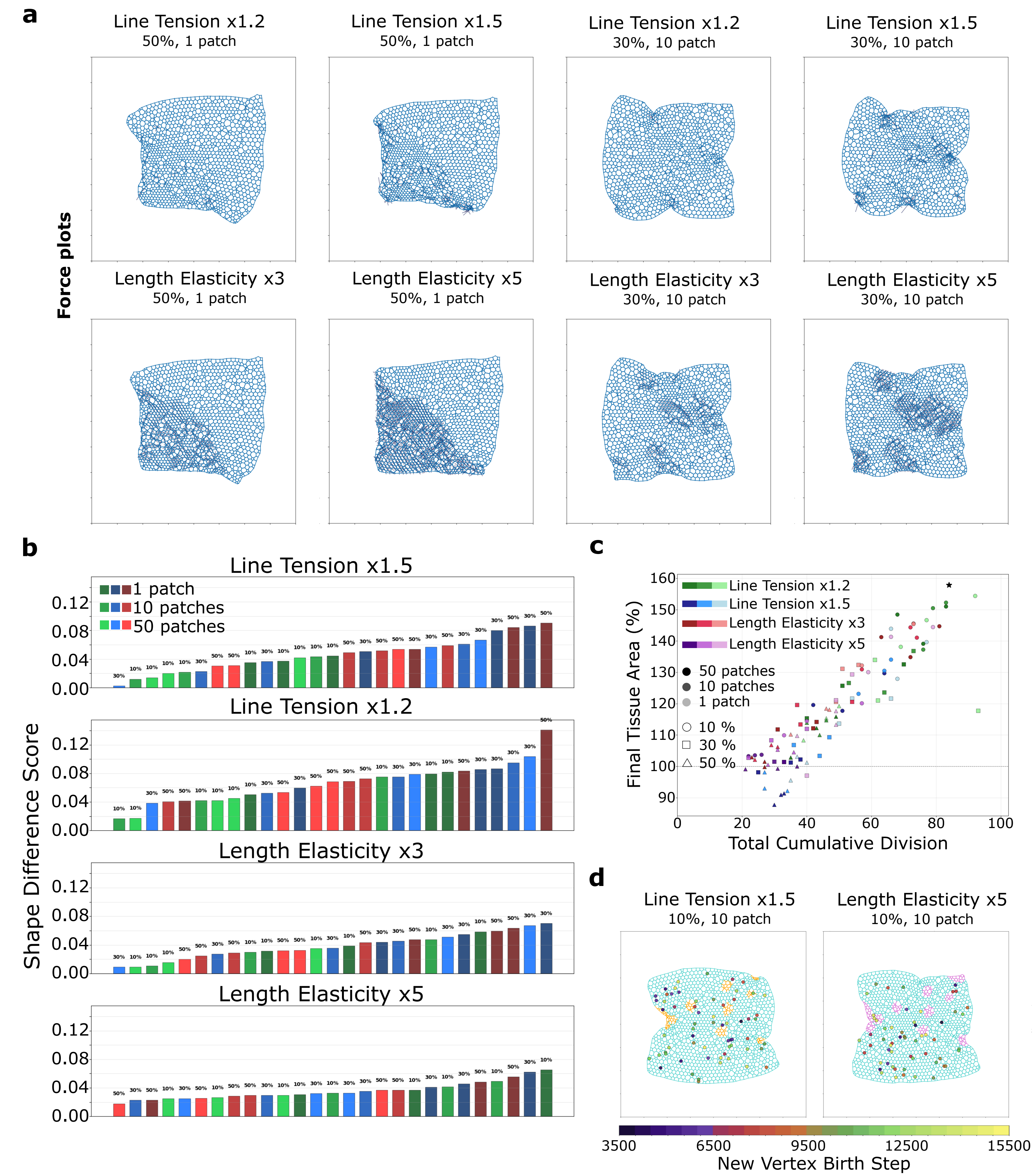


**Extended Data Figure 8 | Characterising tissue shape, area, and FRC division across line tension and length elasticity perturbations. a**, Representative force distributions at the final stage of tissue expansion for simulations with increased line tension (×1.2 and ×1.5) or length elasticity (×3 and ×5). Examples are shown for localised (50% of the network, 1 patch) and dispersed (30% of the network, 10 patches) perturbations. **b**, Shape difference scores for all simulations ranked from lowest to highest shape difference. Bar colour indicates perturbation magnitude (10%, 30% or 50% of the network), and colour shade denotes the spatial distribution (1, 10 or 50 patches). **c**, Relationship between final tissue area and cumulative FRC division events across all simulations. Colours denote perturbation type (line tension ×1.2 / ×1.5, length elasticity ×3 / ×5), point shade indicates the number of perturbed patches (1, 10 or 50), point shape indicates perturbation extent (10%, 30% or 50%), black star represents control simulation. **d**, Representative spatial distribution of FRC divisions after tissue expansion, shown for line tension ×1.5 and length elasticity ×5 (both 10%, 10 patches). Vertices are coloured according to the simulation step at which they were generated. Each condition reflects N = 3 independently initialised simulations. Data in support of Figure 6.
